# Metabolic shift toward ketosis in asocial cavefish increases social-like affinity

**DOI:** 10.1101/2022.05.20.492896

**Authors:** Motoko Iwashita, Amity Tran, Marianne Garcia, Jia Cashon, Devanne Burbano, Vanessa Salgado, Malia Hasegawa, Rhoada Balmilero-Unciano, Kaylah Politan, Miki Wong, Ryan W.Y. Lee, Masato Yoshizawa

## Abstract

Social affinity and collective behavior are nearly ubiquitous in the animal kingdom, but many lineages feature evolutionarily asocial species. These solitary species may have evolved to conserve energy in food-sparse environments. However, the mechanism by which metabolic shifts regulate social affinity is not well investigated. In this study, we used the Mexican tetra (*Astyanax mexicanus*), which features riverine sighted surface (surface fish) and cave-dwelling populations (cavefish), to address the impact of metabolic shifts on asociality and other cave-associated behaviors in cavefish, including repetitive turning, sleeplessness, swimming longer distances, and enhanced foraging behavior. After one month of ketosis-inducing ketogenic diet feeding, asocial cavefish exhibited significantly higher social affinity, whereas social affinity regressed in cavefish fed the standard diet. The ketogenic diet also reduced repetitive turning and swimming in cavefish. No major behavioral shifts were found regarding sleeplessness and foraging behavior, suggesting that other evolved behaviors are not largely regulated by ketosis. We further examined the effects of the ketogenic diet via supplementation with extragenic ketone bodies, revealing that ketone bodies are pivotal molecules positively associated with social affinity. Our study indicated that fish that evolved to be asocial remain capable of exhibiting social affinity under ketosis, possibly linking the seasonal food availability and sociality.

## Introduction

Wild animals experience frequent fasting due to daily, seasonal, and yearly changes in food availability. Physiologically, fasting can increase the secretion of appetite-related hormones (e.g., ghrelin, peptide Y, orexin) and induce a metabolic shift to nutritional ketosis [1]. In terms of behavioral outputs, fasting also induces shifts including boldness in foraging involving risk-taking [2] and a shift from avoiding to approaching prey [3]. Interestingly, fasting also induces non-foraging-related behaviors including aggression towards cohorts [4, 5] and engagement in social dominance [6]. These non-foraging-related behaviors could be evoked by metabolic changes that occur in a state of nutritional ketosis instead of the increased production of appetite-related hormones. However, it is not fully understood whether ketosis itself, in the absence of hunger, drives these non-foraging-related behaviors. Knowledge of such mechanism will open a path to understanding the effects of different dietary intakes according to changing environments, such as switching from glycolysis-inducing carbohydrate-rich diets to ketosis-inducing very low-carbohydrate diets or vice versa.

Recently, the ketosis-inducing ketogenic diet (KD), which contains a high amount of fat, sufficient protein, and a very low amount of carbohydrates, gained popularity among humans because of its neuroprotective and anti-inflammatory effects without impacting appetite-related hormone levels [7–9]. The KD is an effective treatment for refractory seizures, and there is some evidence that it may be beneficial for other nervous system-based disorders, such as Alzheimer’s disease, Parkinson’s disease, and autism [10–13]. Because modern humans evolved to acquire resistance to starvation [14], our body physiology and behavioral tendencies possibly evolved to accommodate drastic metabolic changes. However, the major molecular mechanisms for these possibilities are largely unknown [9, 15]. We were therefore motivated to explore the effects of metabolic shifts, particularly from glycolysis to ketosis, on behavioral outputs such as social affinity using a single species consisting of two morphotypes: typical and starvation-resistant populations.

A suitable model system for this purpose is the Mexican cavefish (*Astyanax mexicanus*). *A. mexicanus* has emerged as a useful experimental platform for diverse aspects of evolution and development, including those with translational relevance to human medicine, such as cataract formation, diabetes, albinism-related syndrome, and insomnia [16–28]. Although there are many parallels in biological phenomena, we strongly agree that the systemic and organ physiologies between humans and this fish species are quite different. Therefore, we do not consider this fish system as an animal model for human disorders but as a platform revealing the genetic and cellular mechanisms that may be conserved among vertebrates and are otherwise difficult to hint at etiologies for similar symptoms. This *A. mexicanus* consists of surface riverine epigean (surface fish) and cave-dwelling hypogean (cavefish) populations. Cavefish diverged from their surface-dwelling relatives 20,000– 200,000 years ago [29, 30], and they have subsequently evolved many distinct morphological and behavioral phenotypes in the food-sparse cave environment, including eye regression/loss, pigment reduction, increased mechanosensory lateral line activity, adherence to vibration stimuli, sleeplessness, hyperactivity, repetitive circling, and reduced social affinity [19, 31–33]. Compared to cavefish, surface fish exhibit typical teleost phenotypes, including typical eyed and pigmented morphologies, no strong adherence to vibration stimuli, nocturnal sleep patterns, and social affinity. Many cavefish traits are believed to have evolved to adapt to food-sparse dark environments. Indeed, wild cavefish are estimated to be exposed to approximately 6 months of food-sparse conditions annually [34], and they are likely to have the ability to withstand starvation via increased fat storage, increased appetites, insulin resistance for fasting [17, 24, 35, 36], slower weight loss during starvation [37], reduced energy-costing circadian activities, and the lack of eyes [38, 39].

Regarding social behavior, cavefish exhibit no detectable schooling behavior [40–42] or hierarchical dominance [43]. By contrast, surface fish school/shoal with cohorts and plastic model fish [40] and exhibit group hierarchical dominance [43]. Because social behaviors in many fish (*e.g.*, zebrafish) are promoted by visual stimuli, blind cavefish might not express social activities because of the absence of visual acuity. However, a recent detailed study illustrated that surface fish exhibit a high level of social-like nearby interactions (one-by-one affinity: social affinity) in the dark, and were promoted by mechanosensory lateral line inputs [33, 41]. In contrast, blind cavefish displayed much lower levels, albeit significant, of nearby interactions than surface fish [33]. Furthermore, cavefish exhibited plasticity in the level of nearby interactions, wherein they increased interaction levels in a familiar environment in comparison with an unfamiliar environment [33]. This observation is similar to those in patients with autism [44, 45] although there is an enormous gap in the complexities of brain functions between humans and fish.

Thus far, similarities between cavefish and patients with autism have been investigated in terms of gene regulation and innate behavior profiles. First, the cavefish gene expression profile is closer to that of patients with autism than to that of other mammalian model systems—a comparison between cavefish and surface fish transcriptomes exhibited the same directional gene expression changes in cavefish observed in the brains of patients with autism (over 58.5% of 3,152 cavefish orthologs). Conversely, other proxy systems (e.g., BTBR mice [classic model for autism] and *shank3* knockout mice) exhibit much less overlap (< 11%) [31, 46, 47]. Second, cavefish’s evolved behaviors, including asociality, repetitive behavior, sleeplessness, higher swimming activity, adherence to a particular vibration stimulus, and higher anxiety-related plasma cortisol levels, are similar to those in patients with autism [31]. Lastly, cavefish and human ancestors are starvation-resistant, and they share some metabolic pathways [14, 17, 24, 37]. These similarities, along with the fact that the ketognic diet (KD) increases socialization in patients with autism [11, 48–50], prompted us to study the effects of ketosis on social affinity in asocial cavefish. Note that we do not consider *A. mexicanus* as an animal model for autism due to substantial differences in their physiologies. What we expect in studies using *A. mexicanus* is that the molecular and cellular responses in ketosis, where the gene expression landscape is the closest to patients with autism, and whose changes are relevant to social behaviors, might provide a hint for the biomedical application that would otherwise be difficult to obtain. This prediction is based on the fact that we have learned so much about human molecular and cellular signaling pathways from fruit flies, which are even phylogenetically more distant organisms than fish [51].

With both the shift in sociality under the seasonal nutrients and the genetic relevance to human disorders in mind, in this study, we assessed the effects of the KD on an evolutionarily asocial cave population of *A. mexicanus*. First, the two-week fasting promoted the level of nearby interaction in asocial cavefish but not in surface fish. Both the two-week fasting and one month of KD feeding reduced the serum glucose-ketone index, whose lower scores (less than 9) indicate ketosis in the body metabolism in human [52], in both surface fish and cavefish. The one-month KD feeding also significantly increased the ketone body concentration in the cavefish brain tissue. Under this feeding regimen, the time-course experiment revealed that one month of KD feeding promoted and sustained the juvenile level of nearby interactions, whereas cavefish fed a control diet (CD) exhibited diminished nearby interactions. KD feeding also reduced repetitive turning and swimming activity. However, the effects of the KD were limited. For example, sleeplessness and high adherence to a particular vibrating stimulus did not show any major changes under the 1-month KD treatment. To reveal the molecular basis of the effects of the KD, we provided supplementation with a major ketone body, beta-hydroxybutyrate (BHB), which promoted social interactions and reduced repetitive turning, covering the major effect of the KD. Finally, we interpreted the possible neural processes influenced by the KD based on affected and unaffected behaviors. According to the study of shared dysregulated genes between cavefish and patients with autism, GO term and KEGG pathway analyses indicated that the dopaminergic system—though less likely the cholinergic, or orexinergic systems—could respond to the KD. However, ketone bodies may have different effects on fish and mammalian physiologies, as discrepancies have been observed in the appetite regulation [53]. Therefore, we need to be careful when interpreting the results of this study in mammals. Nevertheless, *A. mexicanus* provides a unique opportunity to investigate the molecular and genetic responses to ketone bodies in a genetically close biological platform to a human disorder, which will be complementary in understanding deeper etiologies.

Overall, ketosis appears to be capable of significantly shifting the asociality of evolved cavefish toward the surface fish phenotype, providing new insights into the contribution of the diet to social behaviors.

## Methods

### Fish maintenance and rearing in the lab

The *A. mexicanus* surface fish used in this study were the laboratory-raised descendants of original collections created in Balmorhea Springs State Park in Texas in 1999 by Dr. William R. Jeffery. Cavefish used were laboratory-raised descendants originally collected from Cueva de El Pachón (Pachón cavefish) in Tamaulipas, Mexico in 2013 by Dr. Richard Borowsky.

Fish (surface fish and Pachón cave populations) were housed in the University of Hawai‘i at Mānoa *Astyanax* facility with temperatures set at 21 ± 0.5°C for rearing, 24 ± 0.5°C for behavior experiments, and 25 ± 0.5°C for breeding [54, 55]. Lights were maintained on a 12-h:12-h light:dark cycle [54, 55]. For rearing and behavior experiments, the light intensity was maintained at 30–100 Lux. Fish husbandry was performed as previously described [19, 54, 55]. Fish were raised to adulthood and maintained in standard 42-L tanks in a custom-made water-flow tank system. Adult fish were fed a mixed diet to satiation twice daily, starting 3 h after the lights were turned on (Zeitgeber time 3 [ZT3] and ZT9; Zeigler Adult zebrafish irradiated diet, Zeigler Bros, Inc, Gardners, PA; TetraColor Tropical Fish Food Granules, Tetra, Blacksburg, VA, USA; Jumbo Mysis Shrimp, Hikari Sales USA, Inc., Hayward, CA, USA). All fish used in the behavioral experiments were between 2.5 and 5 cm in standard length, fed with live *Artemia* larvae ad libitum, and occasionally supplemented with Dr. Bassleer Biofish Food Fuco (Bassleer Biofish, Herselt, Belgium). They were between 3 and 12 months old. Fish ages were stated for each experiment. All fish care and experimental protocols were approved under IACUC (17-2560) at the University of Hawai‘i at Mānoa.

### Control diet, Fasting, KD, BHB and KE treatments

For the Control diet (CD) feeding, the 4-month-old fish were routinely fed live *Artemia* larvae or the zebrafish standard diet (adult zebrafish irradiated diet: Zeigler Bros, Inc., Gardners, PA, USA) for 3 weeks in the home tank of the experimental fish (Ziplock® containers, S. C. Johnson & Son, Inc., Racine, WI, USA). Fish were grouped into four fish per tank and were fed every morning (ZT 0:00– 3:30) and afternoon (ZT 8:00–12:00). The fish were fed ad libitum during each feeding and any remaining food was removed 1 h after feeding using a pipette. Following standard operating protocols (IACUC (17-2560)), water was changed twice a week, and home tanks were cleaned as usual.

For the fasting experiment, the fish were fasted for 2 weeks (13 full days) before recording the behaviors, while the control fish were fed live *Artemia* larvae. We used two different age groups: 4-6 months old and 11-12 months old. The results from these two age groups were essentially the same. Following standard operating protocols (IACUC (17-2560)), tank water was changed (twice a week), and home tanks were cleaned as usual.

To prepare the ketogenic diet (KD), we used a mixture of a human KD (KetoCal3:1®—nutritionally complete, ketogenic medical food; Nutricia North America, Inc. Gaithersburg, MD, USA) and the zebrafish standard diet (adult zebrafish irradiated diet: Zeigler Bros, Inc., Gardners, PA, USA) in a 5:1 ratio. The gross caloric amounts were 6.99 kcal/g for KetoCal3:1 and 3.89 kcal/g for the zebrafish diet. Regarding the control diet (CD), we used the same KetoCal3:1 and zebrafish irradiated diet mixed at a 1:5 ratio. The KetoCal3:1 powder and ground zebrafish irradiated diet were mixed in the aforementioned ratios and solidified with 1% agar at a final concentration of 20% w/v (2 g of mixture in 10 mL of 1% agar). After solidification, both the KD and CD agar were cut into 3-mm^3^ cubes, and each four-fish group (3-4 months old) was given 1–2 pieces every morning (ZT 0:00–3:30) and afternoon (ZT 8:00–12:00). The fish were fed ad libitum in each feeding and any remaining food was removed 1 h after feeding using a pipette.

To supplement BHB, we used a commercial fish diet (TetraColor Tropical Granules, Tetra, Blacksburg, VA, USA) mixed with BHB (DL-β-Hydroxybutyric acid sodium salt, MilliporeSigma, St. Louis, MO, USA) at a dosage of 10 mg per gram of body weight (10 mg/body g = 78.7 µmol/body g). In detail, fish generally consume 3% of their body weight g per meal. Accordingly, the BHB supplemental diet contained 0.333g/mL of BHB mixed with 0.2 g/mL of the ground Tetra fish diet (20% w/v) in 1% agar. Fish with a body weight of 1 g would consume 30 mg (3%, approximatey 30 µL) of this diet, which contained 10 mg of BHB. The control diet consisted of 20% w/v of the Tetra fish diet in 1% agar. After solidification, both the BHB and control diet agar were cut into 3-mm^3^ cubes, and each group of four-fish group was given 1–2 pieces every morning (ZT 0:00–3:30) and afternoon (ZT 8:00–12:00). The fish were fed ad libitum in each feeding and any remaining food was removed 1 h after feeding using a pipette. Surface fish and cavefish used in this BHB study were 10-11 months old (young adult: 2.0-2.5 cm in the standard length) at the start of the the feeding.

To supplement the ketone ester (KE; D-β-hydroxybutyrate-R 1,3-Butanediol Monoester) [56], we used the adult zebrafish irradiated diet (Zeigler Bros, Inc., Gardners, PA, USA) mixed with KE at a dosage of 3.26 µmol per gram of body weight as used in humans [56]. In detail, similar to supplementing BHB, the KE supplemental diet contained 19.1 mg/mL (=108.5 µmol/mL, MW=176 g/mol) mixed with 0.2 g/mL of ground Zeigler zebrafish diet (20 % w/v) in 1% agar. When a fish with 1g body weight eat 30 mg (3% of body weight) of this diet, it would intake 3.26 µmol of KE. The control diet for KE consisted of 0.2 g/mL of ground Zeigler zebrafish diet, and adjust the calorie content was adjusted with 147.3 mg of sucrose per body g (equivalent to 580.5 calories/body g, the same as 3.26 µmol of KE) in 1% agar. Additionally, since esters have a bitter taste, we standardized it by adding 10 µL/mL of a commercial bitter agent (Symrise, Holzminden, Germany). After the solidification of agar, both the KE and control diet agar were cut into 3-mm^3^ cubes, and each fish group was given 1–2 pieces every morning (ZT 0:00–3:30) and afternoon (ZT 8:00–12:00). The fish were fed ad libitum in each feeding and any remaining food was removed 1 h after feeding using a pipette. Surface fish showed hesitation with the KE and its control diets during the first 2-3 days of this dietary regime by leaving uneaten food, but they consumed all of it afterward. Cavefish never showed uneaten food in this study. Surface fish and cavefish used in this KE study were 6-7 months old (young adult: approximately 2.0 cm in the standard length) when we started the feeding regime.

### Behavior assays

The protocol for social-like nearby interactions was described previously [33]. Briefly, four fish raised in a home tray (15.6 × 15.6 × 5.7 cm^3^ Ziploc containers, S. C. Johnson & Sons, Inc, Racine, WI, USA) were released in a recording arena (49.5 × 24.2 × 6.5 cm^3^) with a water depth of 3 cm on the stage of a custom-made infrared (IR) back-light system within a custom-built black box (75 × 50 × 155 cm, assembled with a polyvinyl chloride pipe frame and covered by shading film). The IR back-light system was composed of bounce lighting of IR LED strips (SMD3528 850 nm strip: LightingWill, Guang Dong, China). The video was recorded at 20 frames/s using VirtualDub2 software (build 44282; http://virtualdub2.com/) with the x264vfw codec for 6 min, and the last 5 min were used for the analysis. After the recording, the fish were returned to the home tray. The X-Y coordinates of each fish were calculated using idTracker software [57] after the video image was processed for background subtraction using ImageJ [33]. These X-Y coordinates were also used for the turning bias analysis. The duration and number of nearby interactions and swimming speed during and after nearby interaction events were calculated using a custom-made MATLAB script (MathWorks Inc., Natick, MA, USA) [33].

The turning bias rate was calculated as 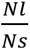, where Ns and Nl represent a smaller (Ns) or larger (Nl) number of left or right turns. This turning bias rate indicates the extent to which fish turning is biased to the left or right, and ranging from “1” (L-R balanced) to infinity (L or R biased). The numbers of left or right turns were calculated as changes in the angles of swimming directions in every five-frame window (0.25 s) as described previously [33]. An automatic calculation of the total number of the left or right turns is implemented in the aforementioned homemade MATLAB script.

Analyses of sleep and swimming distance were described previously [31, 54]. Briefly, fish were recorded in a custom-designed 10.0-L acrylic recording chamber (457.2 × 177.8 × 177.8 mm^3^ and 6.4 mm thick) with opaque partitions that permit five individually housed fish per tank (each individual chamber was 88.9 × 177.8 × 177.8 mm^3^). The recording chamber was illuminated with a custom-designed IR LED source in a light-controlled room on a 12-h:12-h cycle. The room light was turned on at 7:00 am and turned off at 7:00 pm each day. Behavior was recorded for 24 h after overnight (18–20 h) acclimation, beginning 1–2 h after turning the light on (ZT1–2). Videos were recorded at 15 frames/s using a USB webcam with an IR high-pass filter. Videos were captured by VirtualDub2 software with the x264vfw codec and subsequently processed using Ethovision XT (Version 16, Noldus Information Technology, Wageningen, Netherlands). Water temperature was monitored throughout the recordings, and no detectable differences were observed during the light and dark periods (24.0 ± 0.5°C). The visible light during behavior recordings was approximately 30–100 Lux.

The tracking parameters for detection were as follows: the detection was set to “subject brighter than background” and brightness contrast was set from 20 to 255; the current frame weight was set to 15; the video sample rate was set to 15 frames/s; and pixel smoothing was turned off. We monitored sleep activity, and arousal thresholds via protocols previously established for *A. mexicanus* [54]. The X-Y coordinates of each fish were subsequently processed using custom-written Perl (v5.23.0, www.perl.org) and Python scripts (3.8) (https://zenodo.org/record/8137637).

We assayed vibration attraction behavior (VAB) as described previously [54, 58, 59]. Briefly, fish were acclimatized for 4–5 days prior to the assay in a cylindrical assay chamber (325 mL glass dish, 10 cm × 5 cm, VWR, Radnor, PA, USA) filled with conditioned water (pH 6.8–7.0; conductivity 600– 800 μS). During the assays, vibration stimuli were created using a glass rod that vibrated at 40 Hz. The number of approaches to the vibrating rod was video recorded during a 3-min period under infrared illumination. The number of fish approaches within a 1.3-mm radius from the vibrating glass rod was analyzed using the X-Y coordinates of each fish head detected by a trained DeepLabCut model [60])

### Measurement of body

Fish were anesthetized with ice-cold conditioned water (pH: 7.0; conductivity: 700 µS), and their weights were measured after taking pictures with a standard camera (Pentax K-1 DSLR with 35-70 mm zoom lens, Ricoh, Tokyo, Japan). The standard body length and body depth were measured using ImageJ software [61].

### Blood ketone and glucose measurements

All blood samples were collected 2–3 h after feeding. Fish were then deeply anesthetized in ice-cold water, and blood was collected from the tail artery. Blood ketone and glucose levels were measured using either the Abbott Precision Xtra (Abbott Laboratories, Abbott Park, Illinois, USA) or Keto-Mojo GK+ (Keto-Mojo, Napa, California, USA) blood glucose and ketone monitoring system according to the manufacturers’ instructions. The readings of Abbott Precision Xtra were standardized by comparing them with the readings of the same blood sample with Keto-Mojo GK+. Both readings from the Abbott and Keto-Mojo meters showwed a high linear correlation (R^2^ = 0.93, P = 0.000103 and R^2^ = 0.74, P = 0.00565 for glucose and ketone readings, respectively; N = 8). It should be noted that these blood glucose and/or ketone measurements can be affected by hematocrit levels above 65% or below 20%. The hematocrit level of *Astyanax* fish is approximately 30%, in which Pachón cavefish showed slightly higher values (35.56 ± 0.03%, means ± standard error of means) compared to surface fish (28.51 ± 0.03%), as reported by Boggs et al., 2022 [62]. To test the effect of hematocrit level, we diluted the *A. mexicanus* serum 2 times with phosphate-buffered saline (PBS, pH 7.2). The results showed no significant difference in the readings of glucose and ketone bodies between surface fish and cavefish (Additional data 1), suggesting that the differences in the hematocrit levels between the cave and surface fish yielded no detectable effects on the readings of serum glucose and ketone bodies. For ketone measurements in the brain samples, the samples were collected 2-3 h after feeding, following the deep anesthesia in ice-cold water (the same as for the blood samples above). Each brain tissue was carefully collected from the cranium individually and snap-frozen in a 1.5 mL microcentrifuge tube (VWR International, Radnor, PA, USA) chilled with liquid nitrogen. The amount of beta-hydroxybutyrate was measured by following the manufacturer’s instructions of the Ketone Body Assay kit (UV) (MilliporeSigma, Burlington, MA, USA). UV readings were performed by using a BioTek Epoch microplate spectrophotometer (Agilent, Santa Clara, CA, USA).

### Statistical analysis

Regarding the power analysis, we designed our experiments based on a three-way repeated-measures ANOVA with a moderate effect size (f = 0.25), an alpha-error probability of 0.05, and a power of 0.80. The number of groups was eight (surface fish vs. cavefish × non-treated vs. treated × pre-treatment vs. post-treatment). G*Power software [63, 64] estimated that a sample size needed for this experiment was nine per group. Therefore, we aimed to use at least 12 fish in each group for all experiments in this study.

For statistical comparisons of our data, we performed tests including Student’s *t*-test, Wilcoxon signed-rank test, and two- or three-way generalized linear model analyses to compare surface and cavefish, treatment and non-treatment, and pre-treatment and post-treatment. We applied the linear model to non-processed data, the generalized linear model (Poisson family) to discrete data including the blood ketone measurements (multiplied by 10 to apply the Poisson family adjustment for the blood ketone measurements that showed the 0.1 step), and the generalized linear model (Gamma family) to processed data (i.e., turning index). Holm’s post hoc correction was used to detemine which contrasts were significant [65].

Regarding replicates of experiments, we used different individuals for each replicate. Sepcifically, we conducted two biological replicates, using different individuals in each trial. There was no repeated usage of individual fish, except for the time-course experiment. For the experiments measuring sleep, VAB, nearby interactions, and turning bias, we used two biological replicates and confirmed that the averages of the experimental data did not differ significantly. We then merged the data acquired from the two biological replicates and presented it as a single set of results.

The aforementioned calculations were performed using R version 4.0.4 software (packages *car*, *lme4*, and *lmerTest*) [66–68], and all statistical scores are available in Additional data 2 and/or the figure legends.

## Results

From our observations in their wild habitat (the Mexican cave Pachón, Additional data 3), cavefish swam slower and remained near each other more frequently than the lab population. Because the cave environment has a limited diet compared to that of the surface, we predicted that cavefish experience frequent ketosis induced by fasting.

In the 1-year-old surface and cave populations of *A. mexicanus,* 2 weeks of fasting indeed reduced serum glucose levels, leading to a lower glucose ketone index (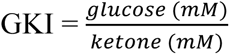; a GKI lower than 9 is considered as ketosis in humans; Additional data 4A-4D; [52, 69]). GKI is proposed as a better indicator of the body ketosis instead of the sole measurement of the ketone bodies [52]. This result indicates that *A. mexicanus* responded to fasting and reduced GKI in a similar manner as mammals, although the serum ketone levels did not significantly increase during this fasting experiment (Additional data 4B). Meanwhile the social-like nearby interactions of cavefish increased (duration and event numbers, Additional data 4E and F; see below for nearby interactions). Although ketosis may be primarily responsible for increasing social interactions, appetite and hormones can also be contributing factors.

Before testing the ketosis-inducing ketogenic diet (consisting of high fat, sufficient protein levels, and very low carbohydrate) to reduce appetite-related behavior, we questioned whether a dietary shift from live *Artemia* larvae (Brine shrimp larvae: BS; standard rearing diet; see Methods) alters the nearby interactions. Feeding Zeigler zebrafish standard diet (control diet: CD) for three weeks did not show any detectable changes in the serum ketone or glucose levels in either surface or cavefish compared to continuous BS-feeding (Additional data 5A and B). Nearby interaction scores were not promoted by this CD feeding; instead, cavefish tended to reduce the scores in both CD and BS-feeding (only detectable in BS-feeding) during 3 weeks of growth (Additional data 5C and D; see below). This reduction of nearby interaction scores was consistent with the following ketogenic diet-feeding study. In summary, the diet shift from BS to CD did not show a detectable effect on serum ketone, glucose, or nearby interaction scores.

Then, to reduce appetite-related behavior, we developed a ketogenic diet (KD) based on a human milk formula (KetoCal3:1® with Zeigler zebrafish standard irradiated diet at a 5:1 weight ratio; Table 1; Materials and Methods).

**Table 1.**
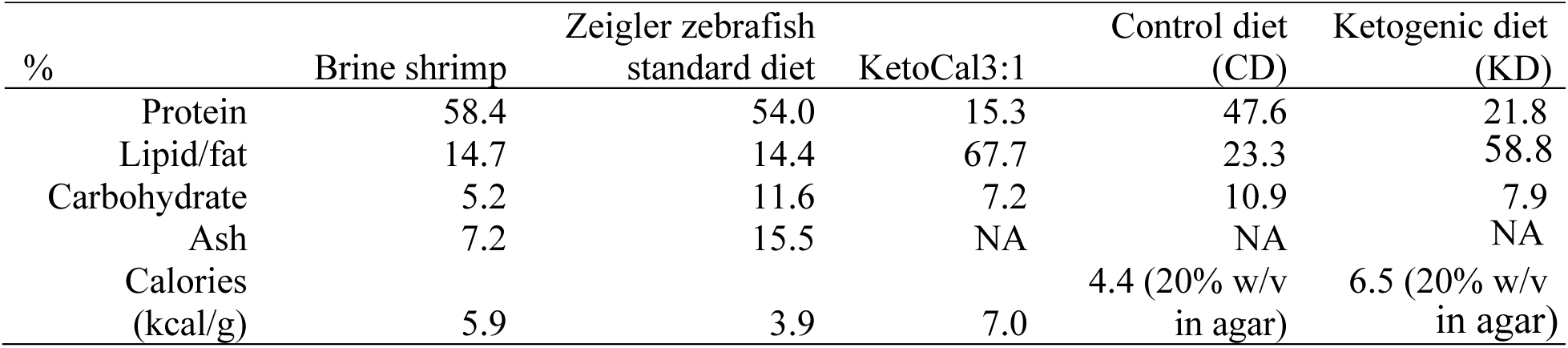
Nutrient composition of each diet used in the study [70, 71]

We then measured the glucose ketone index (GKI) to monitor whether our KD could induce a shift in the balance of ketone bodies and glucose levels after chronic dietary treatment. Three-month-old fish (juvenile–young adult stage) were used in this KD study because the positive effects of KD were more pronounced at the younger stage in human [11], and many adult-type behaviors of cavefish emerge in this stage, including higher adherence to a vibration stimulus (vibration attraction behavior [VAB]) [58], less social affinity, and longer swimming distances compared to surface fish. After 5 weeks of KD feeding, both ketone and glucose concentrations decreased compared to the CD-fed fish (KetoCal3:1 and Zeigler zebrafish diet at a 1:5 weight ratio; Table 1; Figure 1A–C). Surface fish exhibited a significantly higher serum ketone body level than cavefish for both diets (Figure 1B), whereas cavefish exhibited a higher serum glucose level than surface fish (Figure 1C). The GKI was lower in surface fish than in cavefish, and the value was reduced under KD feeding in both surface fish and cavefish compared to that in their CD-fed counterparts (Figure 1D). This result indicats that KD feeding more strongly reduced serum glucose levels than ketone body levels, resulting in a lower GKI in KD-fed fish than in CD-fed fish (Figure 1D). This result suggests that KD feeding could shift the metabolic state from glycolysis toward ketosis. Despite the subtle difference in the serum ketone level between KD-fed and CD-fed cavefish (Figure 1B), a ketone level 8.5 times higher was detected in the brain tissue of KD-fed cavefish than in CD-fed cavefish (Additional file 6; Additional file 2 for the detailed statistical scores). This result implies that the brain cells may efficiently uptake ketones.

**Figure 1.**
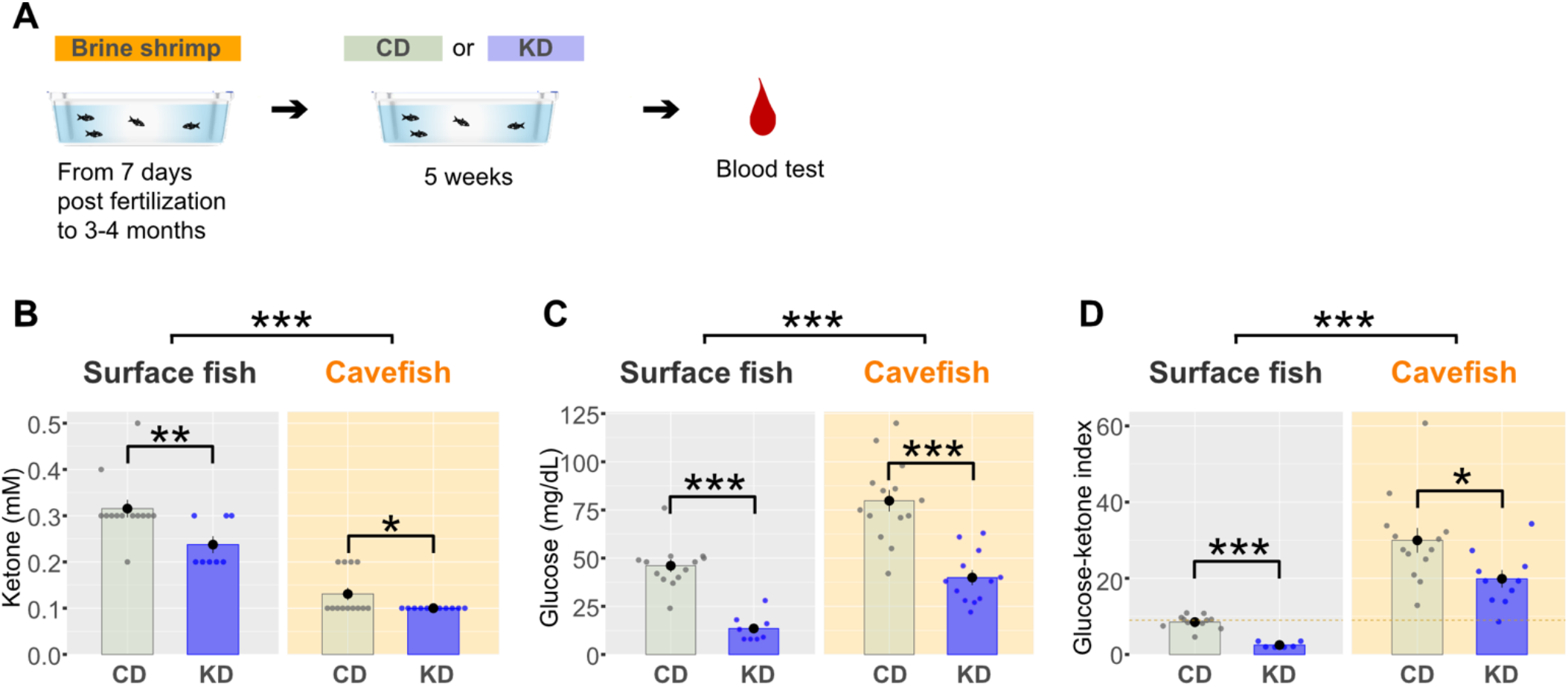
Blood glucose and ketone levels under the control diet (CD) or ketogenic diet (KD). (A) Experimental procedure. After fish were raised for 3–4 months on a brine shrimp larva diet, fish were fed the CD or KD for 5 weeks. Blood glucose and ketone levels were measured after the 5-week period. (B) Blood ketone level (mmol/L). Ketone levels were significantly reduced by KD feeding in both surface fish (SF) and cavefish (CF). Bars represent the data mean and whiskers represent ± standard error of the mean. Dots indicate individual data. The generalized linear model (family = Poisson) followed by post-hoc Holm’s correction was applied for the statistical tests (see Methods and Additional file 2). (C) Blood glucose level (mg/dL). Glucose levels were significantly reduced by KD feeding in both SF and CF. The linear model (family = Gaussian) followed by post-hoc Holm’s correction was applied for the statistical tests. (D) The glucose ketone index (GKI) indicated that the ratio of glucose to ketone was lowered by KD feeding in both SF and CF, suggesting that this diet altered the balance between glucose and ketone. The generalized linear model (family = Gamma) followed by post-hoc Holm’s correction was applied for the statistical tests. SF: N = 13 for CD feeding, N = 8 for KD feeding. CF: N = 13 for CD feeding, N = 11 for KD feeding. *: P < 0.05, **: P < 0.01, ***: P < 0.001. All detailed statistical data are available in Additional file 2.

Regarding this dietary treatment, we first examined its ontogenic (developmental) effects on collective social-like behavior [33]. Many adult behaviors emerge in the transition from juvenile to young adult (adolescent) in 3–4-month-old *A. mexicanus* fish, including foraging behavior, VAB [32, 58], adult-type regulation of sleep (independent from catecholamine) [16, 22, 54, 72], and collective behavior in young adults (under higher Reynold’s number; [33]). Therefore, we therefore investigated the shift in collective behavior in 3–4-month-old fish using our previously reported method [33]. Briefly, it uses criteria based on the vicinity of two fish (≤5 cm) and the duration of nearby interactions (≥4 s) during tracking in a four-fish group (Figure 2B), with criteria defined by permutating the four-fish-group swimming trajectory data 1,000 times [33]. At 3 months old (“Pre-treatment” in Figure 2), surface fish exhibited social-like nearby interactions for 17.0 ± 4.4 s (Figure 2C) and 3.1 ± 0.4 bout number of nearby interactions (Figure 2D) during the 5-min assay. In contrast, cavefish exhibited an approximately 50% shorter interaction duration (8.3 ± 1.5 s; Figure 2C) and a smaller bout number of interactions (1.8 ± 0.3; Figure 2D). To track the effect of the KD treatment, fish were fed the KD for 5 weeks, followed by CD feeding during weeks 6–9 to assess the persistence of the effects of the KD (Figure 2A, C, and D).

**Figure 2.**
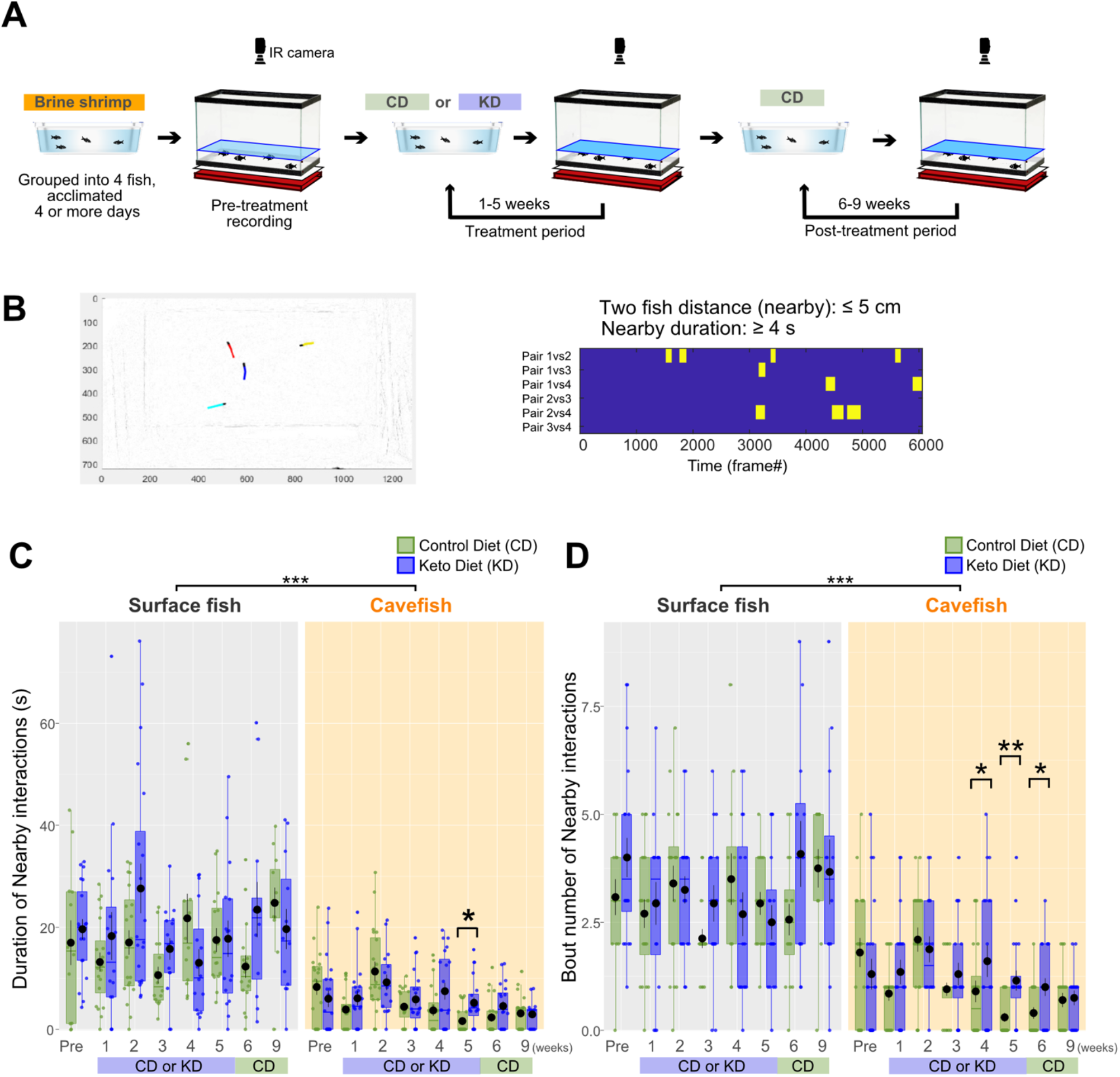
Time-course of nearby interaction changes during 9 weeks of control diet (CD) or ketogenic diet (KD) feeding. (A) Experimental procedure. After rearing fish for 3–4 months on a brine shrimp larva diet, the pre-treatment recording was performed, followed by CD or KD feeding for 5 weeks. Nearby interactions were recorded every week until week 5 of feeding. Subsequently, all groups, including KD-fed fish, were given the CD until week 9. (B) Example of nearby interaction events among surface fish (SF). The left panel presents an example frame of the video, with colored lines indicating the trajectories of individual fish. A red-labeled fish was followed by a blue-labeled fish. Each nearby event that met the detection criteria, namely a distance of ≤ 5 cm between two fish that was maintained for more than 4 s, was counted as a nearby interaction event. The right panel presents an example of the detected events in a raster plot, where each yellow bar indicates a nearby interaction event. Each pair of fish (six pairs among four fish) is presented in the rows. (C) Duration of nearby interactions. Although SF did not exhibit any differences in the duration of nearby interactions (s) between CD (green) and KD (blue) feeding, differences were detected among cavefish (CF) in week 5. However, the nearby interaction duration was indistinguishable from that of the CD group starting in week 6 when the KD was withdrawn from the experimental group. Data are shown in boxplots indicating the 25th, 50th and 75th percentiles in the boxes. The linear mixed-effect model followed by post-hoc Holm’s correction was applied for the statistical tests. (D) Number of nearby interactions. Whereas SF exhibited no differences between CD and KD feeding, differences were observed in CF in weeks 4–6. After the KD was withdrawn in week 6, the number of events decreased to the level observed with CD feeding. Data are presented as boxplots indicating the 25th, 50th, and 75th percentiles. The generalized linear model (family = Poisson) followed by post-hoc Holm’s correction was applied for the statistical tests. Dots indicate individual data. N = 20 for each group. *: P < 0.05, **: P < 0.01, ***: P < 0.001. All detailed statistical data are available in Additional file 2.

The nearby interactions of surface fish did not differ between CD and KD feeding (Figure 2C and D). In contrast, the nearby interactions of cavefish were significantly decreased by CD feeding compared to the effects of KD feeding in weeks 4 and 5 (Figure 2C and D), and interactions remained depressed through week 9 with CD feeding. However, the effect of the KD diet on nearby interactions did not persist. After KD deprivation and CD feeding, the nearby interactions of KD-fed cavefish were indistinguishable from those of CD-fed fish (6–9 weeks, Figure 2C, and D), suggesting that KD has a promotive/supportive effect on collective behavior in genetically asocial cavefish.

To support the finding of KD-promoted nearby interactions with alternative data, we explored the swimming speed profile, which is a supportive indicator of fish movement in the vicinity of each other. Fish are more likely to have an opportunity to express affinity towards each other at a slower swimming speed, although it may not be as obvious at a very slow speed as below 3 cm/s [33]. Consistent with the former report, CD-fed surface fish moved slower during nearby interactions than during the non-nearby interaction period, and a similar speed profile was also observed in KD-fed surface fish [33] (“5 weeks”, Figure 3A). There was no detectable difference in swimming speed profiles between CD and KD feeding (“5 weeks”, Figure 3A). Similarly, KD-fed cavefish swam slower during the nearby interaction period than during the non-nearby interaction period (“5 weeks” Figure 3B). The overall swimming speed was also slower in the KD group than in the CD group (“5 weeks” Figure 3B). These findings indicate that KD-fed cavefish exhibited more social-like nearby interactions with a similar speed profile to surface fish. The effect of KD feeding on the ontogeny of swimming speed/distance showed a somewhat surprising result. We tracked the total swimming distance within 5 min from pre-treatment to week 9 of feeding (Additional file 7). KD-fed cavefish exhibited a significantly shorter swimming distance (indicating slower swimming speeds) from the first week of feeding (Additional file 7), which was much earlier than when the higher level of nearby interactions emerged (weeks 4–5). This result suggests that KD feeding induced calmer swimming in cavefish within a week of the treatment, although a slower speed itself is not sufficient to induce nearby interactions.

**Figure 3.**
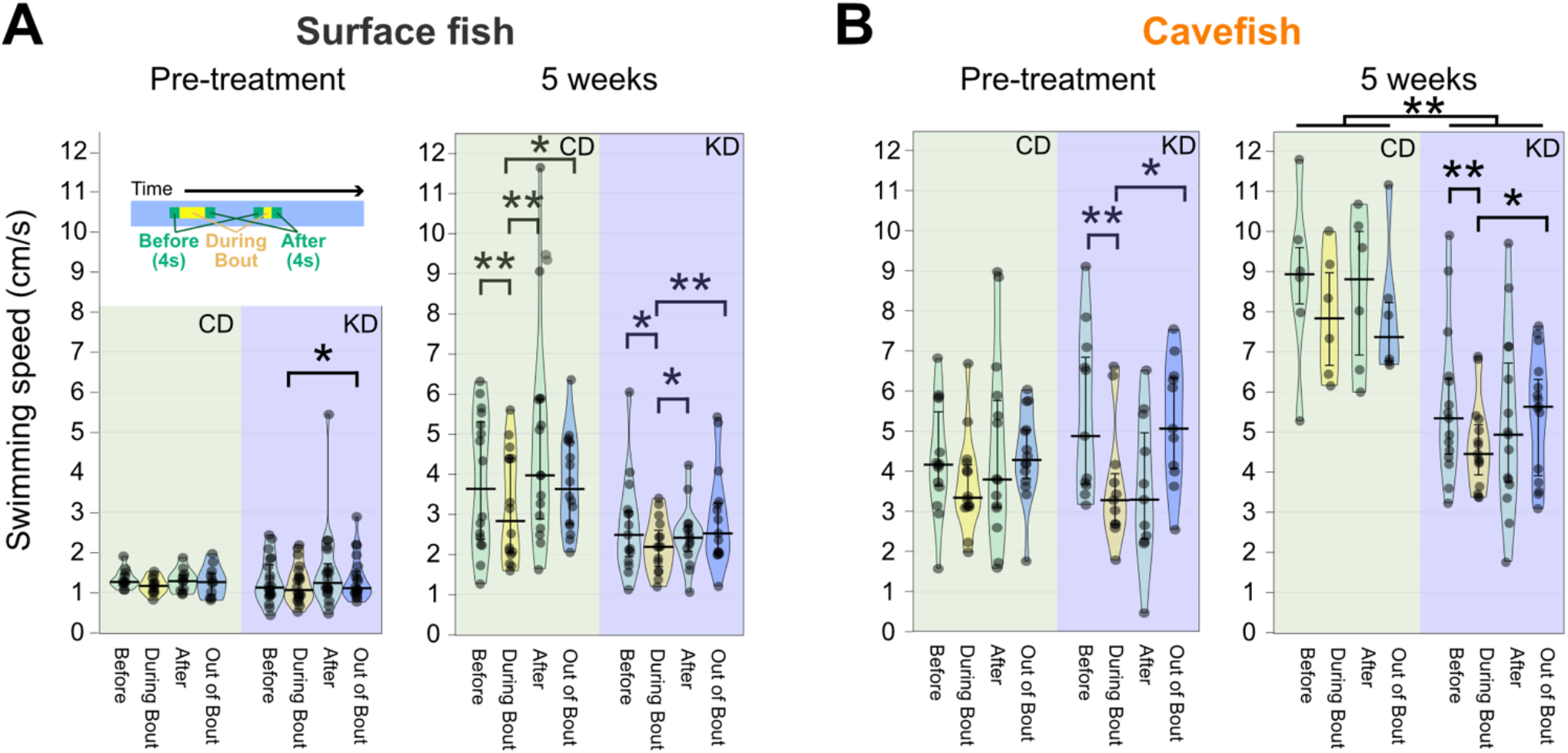
Ketogenic diet (KD) feeding induced surface fish (SF)-like speed profiles during nearby interactions in cavefish (CF). Changes in swimming speed before, during, and after nearby interaction events in SF (A) and CF (B). The mean swimming speeds were plotted for: (i) 4 s before the nearby interaction event, (ii) during the event, (iii) during 4 s after the event, and (iv) during the out-of-event period (see the top-left inset of A). (A) Swimming speed was reduced during nearby interactions in SF in both the CD and KD groups. This profile was clearer in the fifth week (right panel). The linear mixed-effect model followed by post-hoc Holm’s correction was applied for the statistical tests. (B) Swimming speed was reduced during nearby interactions only in the KD group in week 5 (right panel). The bars indicate the 25^th^, medians, and 75^th^ percentiles of the data points. The different speed profiles between the CD and KD groups in the Pre-treatment are due to the naturalistic standing variation in *A. mexicanus* system. The linear mixed-effect model followed by post-hoc Holm’s correction was applied for the statistical tests. Dots indicate individual data. SF: N = 11 for CD, N = 20 for KD. CF: N = 16 for CD, N = 15 for KD. *: P < 0.05, **: P < 0.01, ***: P < 0.001. All detailed statistical data are available in Additional file 2.

Repetitive turning is frequently observed in an antagonistic relationship with nearby interactions in cavefish and mammals [33, 73, 74]. That is, individuals with few nearby interactions frequently exhibit a high level of turning bias or “repetitive turning.” Accordingly, CD-fed cavefish with few nearby interactions exhibited a significantly higher turning bias than KD-fed cavefish (Figure 4A, B). KD-fed cavefish displayed a comparable level of balanced turning as surface fish (close to a score of “1” in Figure 4B). In summary, these results suggest that KD feeding could reduce repetitive turning while inducing longer nearby interactions in cavefish.

**Figure 4.**
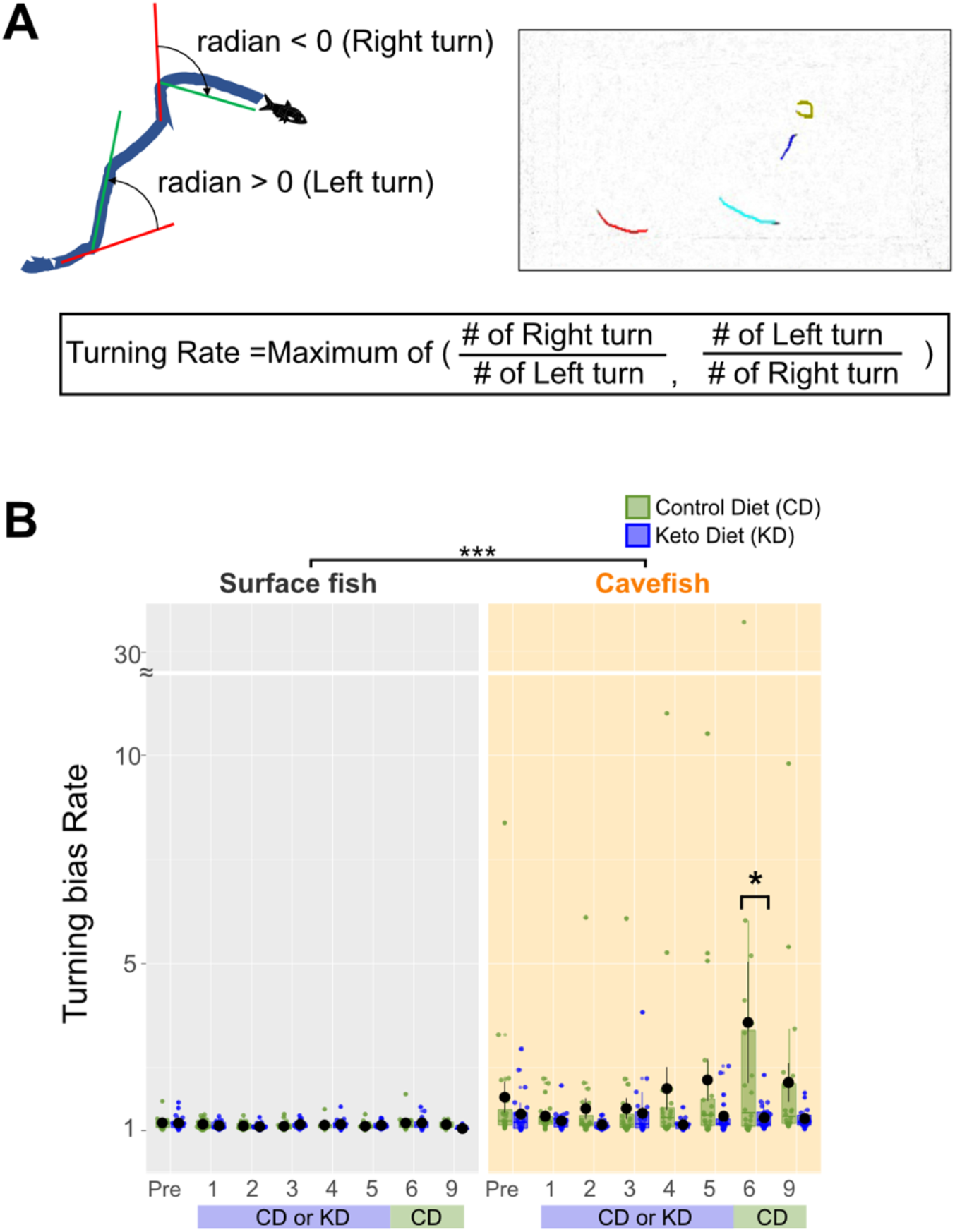
Biased turning was attenuated by the ketogenic diet (KD). (A) Diagram and the calculation formula for the turning bias index. The changes in the left or right traveling directions were calculated every five frames (every 0.25 s) across all trajectories and expressed as radians. Positive radian values represent left (anticlockwise) turning, and negative values indicate right turning. The ratio between the numbers of clockwise and anticlockwise turns was used as the turning rate (1–infinity, positive value). (B) Turning biases of surface fish (left) and cavefish (right). There was no difference between CD and KD feeding in surface fish, whereas the turning index in CD-fed cavefish was larger than in KD-fed cavefish (see week 6). The generalized linear model followed by post-hoc Holm’s correction was applied for the statistical tests. Bars represent the data mean and whiskers represent ± standard error of the mean. Dots indicate individual data. N = 20 for all groups. *: P < 0.05, **: P < 0.01, ***: P < 0.001. All detailed statistical data are available in Additional file 2.

Recording behavior each week (Figure 2) may yield a confounding factor, such as fish remembering the recording environment and behaving differently than naive fish. To clarify whether our results captured the genuine effects of KD feeding, we repeated the 4–5-week dietary treatment in a new set of fish (Additional file 8). As observed in Figure 2, surface fish did not exhibit a detectable change in the duration and number of nearby interactions between CD and KD feeding (Additional file 8A, B). In contrast, CD-fed cavefish displayed fewer nearby interactions, whereas the level of nearby interactions was retained in KD-fed cavefish, resulting in a higher level of nearby interactions in KD-fed cavefish (Additional file 8A, B). In this repeated experiment, the results for repetitive turning were also similar to those in the previous experiment; specifically, CD-fed cavefish displayed a high level of turning bias, whereas KD-fed cavefish exhibited balanced turning (Additional file 8D).

We then explored other changes induced by KD feeding, including changes in sleep, 24-h swimming distance, and adherence to a vibrating stimulus, which are distinct between surface fish and cavefish. Cavefish exhibit reduced sleep duration and swim almost all day, perhaps to find nutrients in the food-sparse environment [16, 20, 22, 54]. After 5 weeks of dietary treatment on the 3–4-month-old fish, both surface fish and cavefish exhibited shorter sleep duration than observed before treatment, regardless of the diet (Figure 5A, particularly at night), suggesting that growth between 3–4 and 4–5 months old exerted a negative effect on the sleep duration. However, there was no detectable difference between CD and KD feeding.

**Figure 5.**
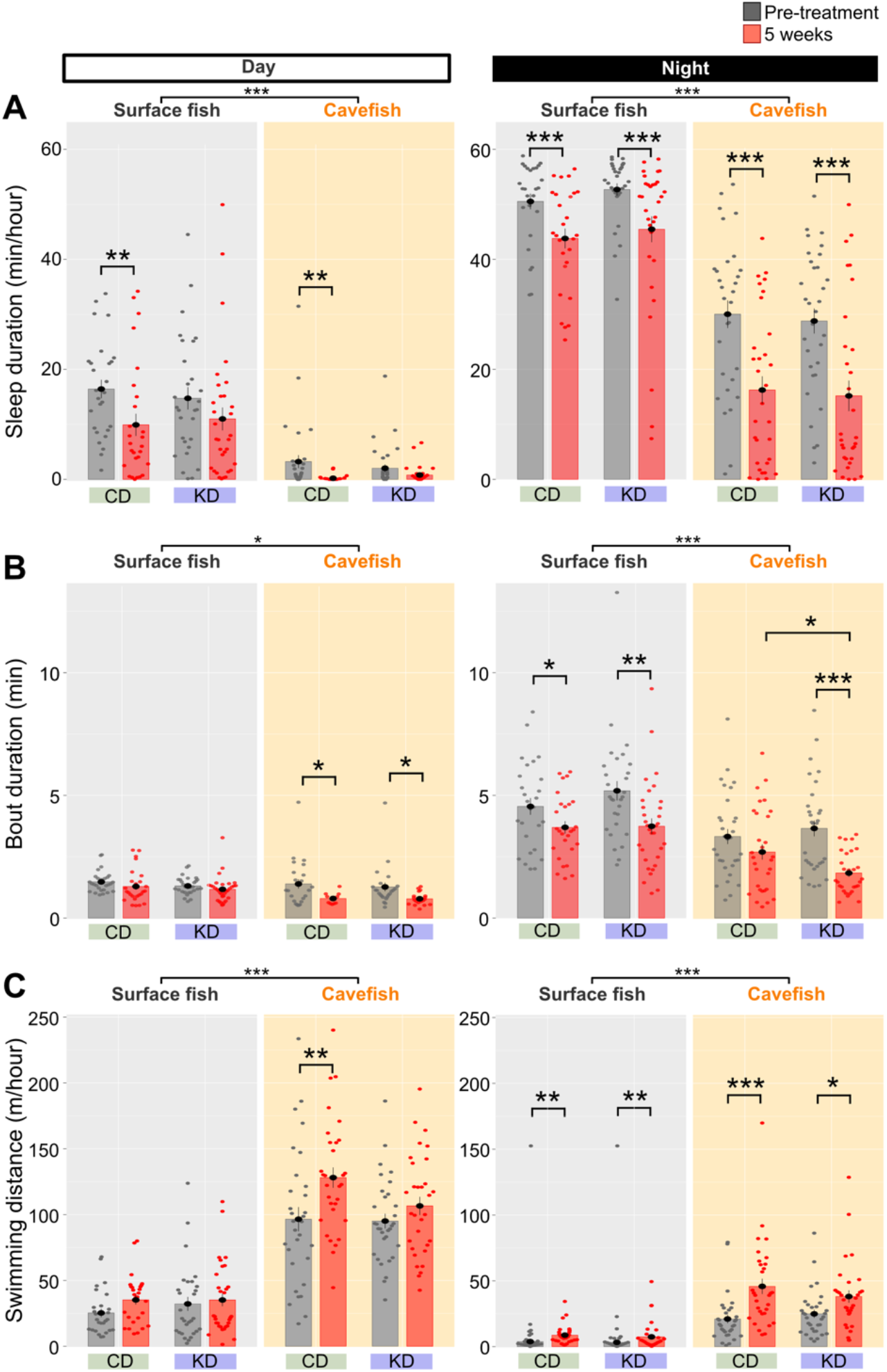
Day and night sleeping durations and swimming distances were not altered by ketogenic diet (KD) feeding. (A) Sleep duration (min/h) during the day (left) and night (right). During 5 weeks of growth, the sleep duration decreased in both surface fish and cavefish regardless of the diet (particularly during night). The linear mixed-effect model followed by post-hoc Holm’s correction was applied for the statistical tests. (B) Average sleep-bout duration (min/10 min bin) during the day (left) and night (right). During 5 weeks of growth, the sleep bout duration was lower in surface fish under both dietary conditions and in KD-fed cavefish (night). The linear mixed-effect model followed by post-hoc Holm’s correction was applied for the statistical tests. (C) Swimming distance during the day (left) and night (right). Cavefish fed the control diet (CD) exhibited a longer swimming distance during the day and night. Conversely, both surface fish fed either diet and cavefish fed the KD exhibited a significantly increased swimming distance only at night. The linear mixed-effect model followed by post-hoc Holm’s correction was applied for the statistical tests. Bars represent the data mean and whiskers represent ± standard error of the mean. Dots indicate individual data. Surface fish: N = 28 for CD, N = 32 for KD. Cavefish: N = 28 for CD, N = 32 for KD. *: P < 0.05, **: P < 0.01, ***: P < 0.001. All detailed statistical data are available in Additional file 2.

Animals’ sleep is usually fragmented, involving repeated sleep/awake cycles during the night (diurnal animals) or day (nocturnal animals) [75]. Then, the structure and regulation of sleep are typically analyzed according to the average duration and the number of events (bouts). Our detailed sleep analysis illustrated that KD-fed cavefish displayed shorter sleep bout duration during the night than CD-fed cavefish (Figure 5B). However, the number of sleep bouts did not differ between CD and KD feeding (5 weeks; Additional file 9). Overall, the sleep phenotype showed a subtle change by KD feeding, as cavefish exhibited shortened sleep duration.

Sleep duration is negatively correlated with the 24-h swimming distance [54]. Cavefish displayed overall higher activity, which was consistent with previous findings [20, 54], and consistent with the findings of longer swimming distances in the nearby interaction assay (Figure 5C). CD-fed cavefish swam longer distances after the 5-week treatment, but KD-fed cavefish did not show a detectable change in it after the treatment (during the day period, Figure 5C). Surface fish, in contrast, did not exhibit a detectable difference in swimming distance between KD and CD feeding. Overall, the KD treatment induced subtle changes in sleep-associated behaviors in both surface and cavefish.

In general, the KD is assumed to induce ketosis without increasing appetite. We then checked the shift in foraging behavior under KD feeding. Cavefish evolutionarily exhibit increased foraging behavior, which can be quantified with vibration attraction behavior (VAB), in which fish adhere to a particular vibration stimulus (35–40 Hz) in the dark [58]. VAB is advantageous for prey capture in the dark. Cavefish and surface fish did not exhibit a detectable difference in VAB between CD and KD feeding, whereas VAB was significantly increased during 1 month of growth (pre-treatment vs. 5 weeks; Additional file 10A). In summary, the VAB analysis indicated that KD feeding did not significantly increase or decrease foraging behavior.

Although the KD diet induced significant changes in some behavioral outputs, it suppressed growth during treatment. The average weights of KD-fed surface fish and cavefish were 55.5 % and 69.9 % of those in their CD-fed counterparts, respectively (5 weeks; Figure 6B). The standard length of KD-fed surface fish was also significantly reduced (5 weeks; Figure 6A).

**Figure 6.**
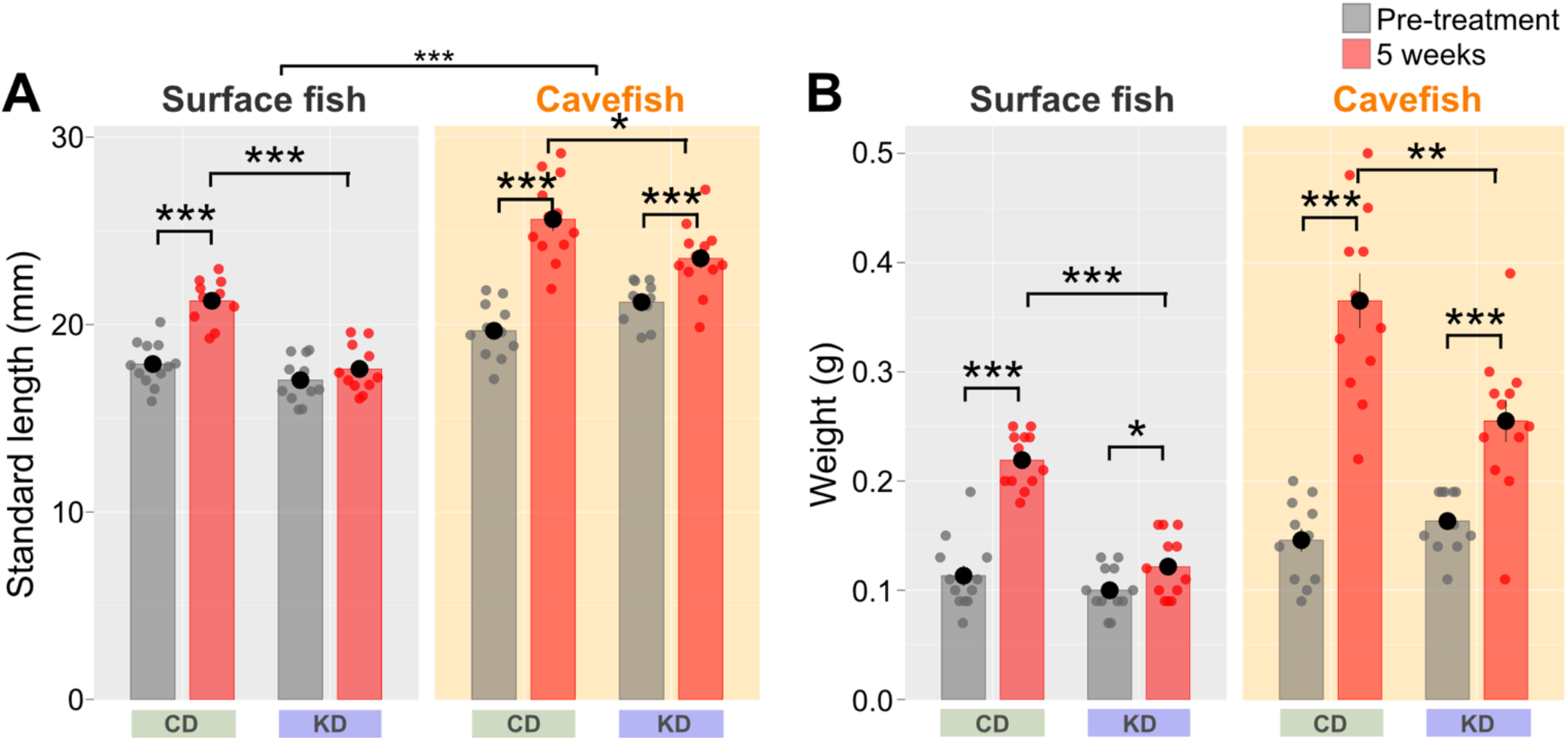
Body length and weight under control diet (CD) or ketogenic diet (KD) feeding. (A) Standard length (cm). KD-fed surface fish and cavefish were significantly smaller than their CD-fed counterparts. The linear mixed-effect model followed by post-hoc Holm’s correction was applied for the statistical tests. (B) Body weight (g). KD-fed surface fish and cavefish weighed less than their CD-fed counterparts. The linear mixed-effect model followed by post-hoc Holm’s correction was applied for the statistical tests. Data are presented as the mean ± standard error of the mean. Dots indicate individual data. Surface fish: N = 28 for CD, N = 32 for KD. Cavefish: N = 28 for CD, N = 32 for KD. *: P < 0.05, **: P < 0.01, ***: P < 0.001. All detailed statistical data are available in Additional file 2.

Are these behavioral and growth changes induced by ketosis? The KD contains high amounts of fat, sufficient levels of proteins, and a minimum amount of carbohydrates. This question motivated us to test the molecular basis of the effects of KD feeding by supplementing major ketosis metabolites, ketone bodies, to the standard diet.

In humans, KD feeding induces ketosis, in which the liver releases beta-hydroxybutyrate (BHB) and acetoacetate via beta-oxidation of fat [76]. Instead of supplying a massive amount of fat using the KD, BHB might be responsible for the majority of effects observed after KD feeding. With this idea, the ketone ester (D-β-hydroxybutyrate-R 1,3-Butanediol Monoester; delta-G® [56]) was provided as a supplement to both surface fish and cavefish for 5 weeks. The ketone ester (KE) was expected to undergo complete hydrolysis by the gut esterases, resulting in two BHB molecules (and acetoacetate)[56]. It does not contain any salt ions, unlike the sodium or potassium salt forms of BHB, nor does it has the racemic L-form; where only the D-form is considered to be biologically active [77]. Since we were unsure whether gut esterases were available in juvenile-adolescent fish at 3 months old, we used 6-7 months old fish that have a mature gut system but are in the young adult stage. The results indicated that the KE supplementation significantly reduced the serum GKI (Additional file 11), while promoting nearby interactions in cavefish (Figure 7A and B). Swimming distance was slightly reduced in cavefish (Figure 7C). Turning bias was not reduced by KE supplementation in cavefish (Figure 7D). There was no detectable difference between CD and KE supplemental diets in sleep duration or VAB (Additional file 12A and 12B, respectively).

**Figure 7.**
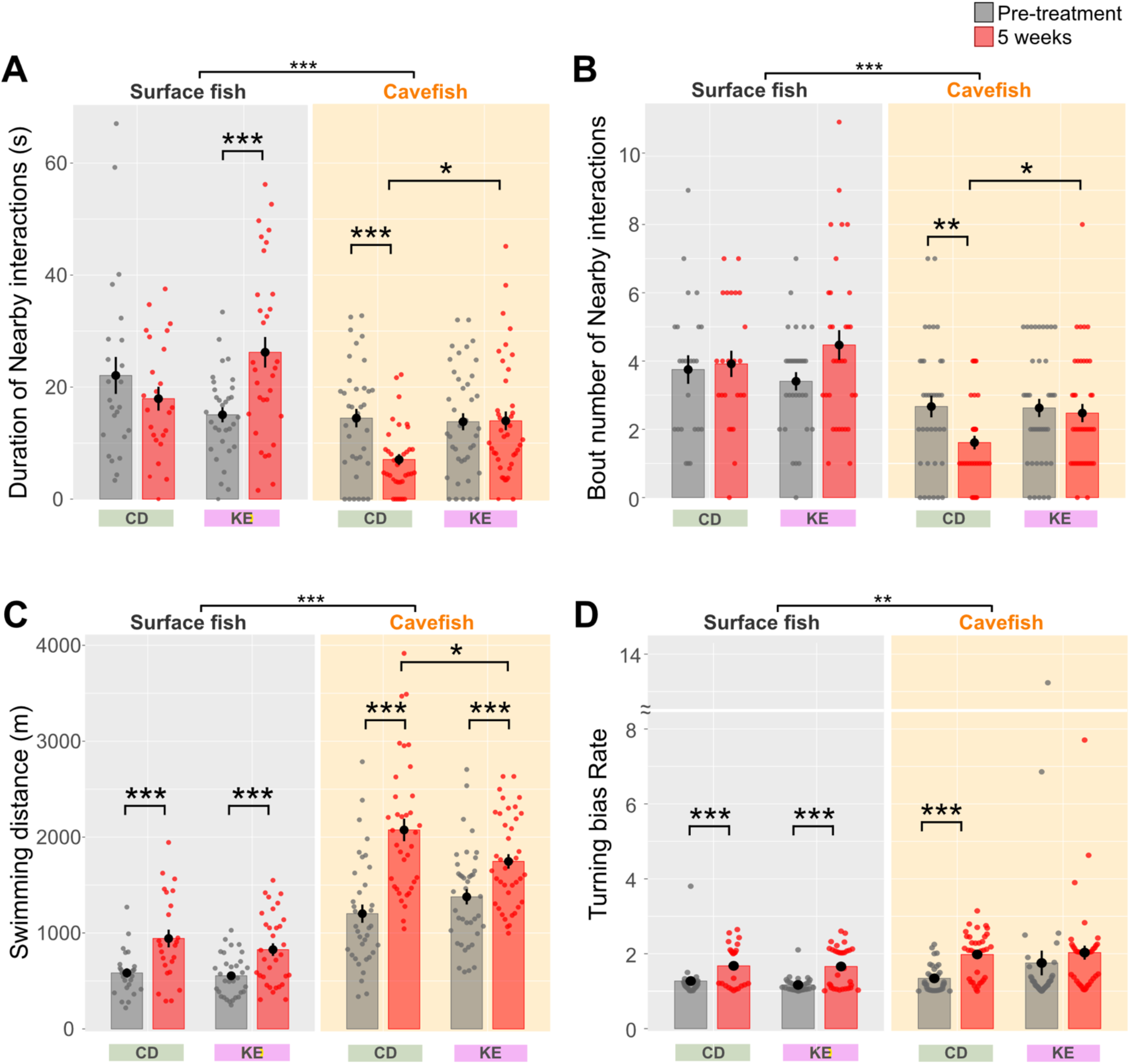
Nearby interactions and other behaviors under control diet (CD) or ketone ester– supplemented diet (KE) feeding. (A) Duration of nearby interactions (s). After 5 weeks, the duration of nearby interactions increased in KE-treated surface fish and cavefish. The linear mixed-effect model followed by post-hoc Holm’s correction was applied for the statistical tests. (B) Number of nearby interaction events. The number of nearby interactions was promoted in KE-treated cavefish. The generalized linear model (family = Poisson) followed by post-hoc Holm’s correction was applied for the statistical tests. (C) Swimming distance. KE-treated cavefish exhibited a slight but significant decrease in swimming distance compared to CD-treated cavefish in a 5-min assay. The linear mixed-effect model followed by post-hoc Holm’s correction was applied for the statistical tests. (D) Turning bias ratio. No significant difference was detected between the CD and KE groups. The generalized linear model (family = Gamma) followed by post-hoc Holm’s correction was applied for the statistical tests. Data are presented as the mean ± standard error of the mean. Dots indicate individual data. N = 20 for all groups. *: P < 0.05, **: P < 0.01, ***: P < 0.001. All detailed statistical data are available in Additional file 2.

Interestingly, the body growth of KE-treated surface and cavefish was not significantly different from that of the control diet (CD), suggesting that the KE molecule (and therefore, the BHB molecule) did not have a detectable negative effect on growth (Additional file 12C and D) (cf. Figure 6 in KD).

We also tested the supplemental feeding of the BHB salt form (sodium salt form of racemic BHB: 50% L-form and 50% D-form). We used 11-12 months-old fish in this study since the younger fish seemed to suffer from the high-salt-containing diet. The 4-week feeding result was essentially the same as the KE supplemented diet feeding: the BHB salt supplemental diet significantly reduced GKI in the serum of surface and cavefish (Additional file 13), while promoting nearby interactions in cavefish (Additional file 14A and B) but reduced the duration of nearby interactions in surface fish (Additional file 14A). No major change in response to the BHB feeding was detected in, swimming distance (Additional file 14C), turning bias (Additional file 14D), sleep (Additional file 15A) and VAB (Additional file 15B) in cavefish, while the BHB salt reduced growth (standard length and weight) in surface fish (Additional file 15C and D). In contrast, cavefish did not show any detectable negative effects on growth under the BHB salt supplemental feeding (Additional file 15C and D).

In summary, BHB (KE and BHB salt) treatment encompassed the effect of the KD treatment—promoting social interactions. BHB, particularly KE, had a no-detectable negative effect on growth. These facts suggest that ketone bodies can be responsible factors for the positive effects on social behaviors of KD feeding. BHB treatment also indicated that older-age cavefish (6-7 months or 11-12 months old) were still capable of responding to ketone bodies, not only younger age groups (3-4 months old).

## Discussion

In this study, we examined the behavioral shifts induced by KD feeding and supplementation with ketone bodies. Ketosis (high metabolic usage of ketone bodies comapring with glucose) is expected to occur frequently in wild animals due to a failure to find food (fasting) or a reduced carbohydrate inputs/synthesis (available nutrients). Even in asocial species that have evolutionarily reduced sociality levels, certain levels of social interaction can still be crucial for mating. Under KD feeding, cavefish maintained their juvenile level of nearby interactions until the treatment ended (5 weeks). Subsequently, within 1 month after stopping KD feeding, nearby interactions were reduced to an indistinguishable level from the control group. Surface fish exhibited a higher number of nearby interactions than cavefish, and no detectable difference was observed in nearby interaction levels between CD- and KD-fed surface fish. KD feeding also effectively reduced repetitive turning in cavefish, whereas CD-fed cavefish exhibited a high level of repetitive turning. There were no major changes in sleep duration and foraging behavior (VAB) after 1 month of KD feeding. These patterns in behaviors and growth were consistent across two replicated experiments (social affinity and repetitive turning), supporting a scientific rigor of the observed effects under KD feeding. Additionally, the diet shift from live brine shrimp (the standard diet for juveniles) to the standard zebrafish pellet diet (used as the control diet in this study) did not yield any detectable changes in behaviors, serum glucose, and ketone levels, further supporting the effect of KD feeding. Finally, the major KD metabolite, BHB, could account for KD’s positive effect on nearby interactions, indicating that the ketone bodies play a pivotal role in this treatment.

### Effects of the KD on blood ketone levels and body growth

During 4–5 weeks of KD feeding, blood ketone and glucose levels were reduced compared to the effects of the CD in both surface fish and cavefish, contradicting our expectation of higher serum ketone levels in the KD group. However, the GKI [52] was significantly lower under KD feeding than under CD feeding. These significant changes in GKI in both surface fish and cavefish suggest that the metabolic balance shifted toward the ketosis side due to KD feeding. Indeed we detected the higher ketone concentration in cavefish brain tissues under KD feeding compared to CD feeding. In general, cavefish had a higher GKI than surface fish under both diets, indicating that the cavefish physiology was constitutively biased toward glycolysis. For example, blood glucose levels in cavefish under KD feeding were similar to those in surface fish under CD feeding, while cavefish had ∼2.5-fold lower ketone levels than surface fish under CD feeding, resulting in a higher GKI even under KD feeding (Fig. 1).

KD feeding for 4–5 weeks also resulted in slowed body growth. This growth retardation has been observed in patients with epilepsy who were chronically fed a KD [49, 78], and these results were consistent with our observations in KD-fed fish. This study found that a ketone body (ketone ester) did not suppress body growth while increased the social-like activities. The detailed molecular/physiological mechanisms by which ketosis affects behaviors are at the early stage of investigations (see murine resaerches [15, 79, 80]). Our studies using ketone ester and BHB sodium salt provided an alternative starting point to unravel the mechanisms underlying KD-associated phenotypes (see below).

### Effects of ketones in the TCA cycle and epigenetics in the brain

In mammals, KD feeding induces a “starvation”-like state, causing the liver to release ketone bodies into the bloodstream. BHB is the major ketone body produced by the liver through beta-oxidation. The gut epithelia also absorb and circulate ketone bodies from the diet and/or gut microbiota. Both liver-derived and gut-derived ketone bodies can cross the blood–brain barrier and serve two functions: (i) inhibiting histone deacetylase, which influences epigenetic regulation and induces gene expression in neurons; and (ii) acting as a general energy source that is converted into acetyl-CoA to fuel the aerobic TCA cycle in neurons. Both pathways have the potential to alter brain function. The fact that cavefish can easily tolerate high blood glucose levels that would paralyze surface fish [24], and the upregulated Wnt signaling in cavefish, potentially resulting in high glycolytic activity as observed in humans [31, 81], support the aforementioned hypothesis that cavefish exhibit high blood glucose levels and mainly generate energy via glycolysis. Additionally, ketone bodies can promote behavioral shifts by changing the epigenetic state through histone deacetylase (HDAC) inhibition [82, 83]. Inhibition of HDAC increases gene expression in general. This possibility may lead to a positive effect in cavefish neurons due to the fact that cavefish have more downregulated genes (2,913 genes, log_2_ < −1.0) than upregulated genes (1,643 genes, log_2_ > 1.0) in the transcriptome at 72 h postfertilization [31, 84]. Furthermore, more methylated loci are found in the eye genes of cavefish than in surface fish, which could also occur in other tissues including the brain regions [85], and most of these methylated gene loci were downregulated. The brains of patients with autism are also expected to be hypermethylated, resulting in a transcription-less condition [86]. Therefore, these two pathways, namely metabolism and epigenetics, are highlighted as possible targets of ketone bodies during behavioral shifts under ketosis. Future research should address these possibilities to clarify the metabolism-based evolution of behavior (*cf.* [15])

### Ontogeny of nearby interactions and the KD

In this study, 3–4-months-old cavefish exhibited a detectable level of nearby interactions (social affinity), which decayed under CD feeding. Interestingly, KD-fed cavefish and surface fish fed either diet maintained a similar level of nearby interactions during the 5 weeks of dietary feeding. The reduction of nearby interactions in CD-treated cavefish can be explained by (1) quicker exhaustion under CD feeding (aerobic ketosis produces more adenosine triphosphate than anaerobic glycolysis), (2) greater anxiety in the recording environment [33], and (3) less social motivation. The first explanation is unlikely because CD-fed cavefish swam at comparable or longer distances than KD-fed cavefish (e.g., Additional file 7 and 8C). The higher level of anxiety could explain the findings because cavefish exhibited increased repetitive turning, which is related to higher anxiety in mammals [74]. In addition, prior rerearch has shown that cavefish displayed fewer nearby interactions in an anxiety-associated unfamiliar environment [33]. In the future, the anxiety level should be monitored using plasma cortisol levels [87]. Less motivation regarding social affinity is also a possible cause, and this possibility can be assessed by examining neural activities in social decision-making networks, including the preoptic area, nucleus accumbens, and striatum [88, 89]. Explanations (2) and (3) are not mutually exclusive, and co-occurrence is possible. These possibilities will be assessed in our future study.

### Possible target system for ketosis

Under KD feeding and BHB treatment, we observed an increased in social affinity and a reduction in repetitive turning. However, we did not detect major changes in sleep and VAB.

Studies regarding neurotransmitters and their associated behaviors have revealed tight associations between them, such as the dopaminergic system being associated with social and repetitive behavior, and the cholinergic/orexinergic/histaminergic systems playing a major role in sleep regulation (Additional file 16). The behavioral phenotypes in this study highlighted the possible involvement of the dopaminergic system but less involvement of others (i.e., the serotonergic, cholinergic, orexin/hypocretinergic, histaminergic, or adrenergic system). The dopaminergic and other social/repetitive behavior-associated pathways were also highlighted in the GO term/KEGG pathway analysis using genes showing the same directional expression changes (upregulation or downregulation) in patients with autism versus neurotypical individuals [90] and cavefish versus surface fish at 72 hrs postfertilization [31, 91, 92] (Additional file 17). The shared same directional expression genes are enriched in the synaptic vesicle cycle, long-term depression/potentiation, dopaminergic/serotonergic synapses, and oxytocin-signaling pathway (Additional file 18). The underlying reason why ketosis or ketone bodies have a stronger effect on the dopaminergic system than on the other systems is undetermined, calling for further investigation. However, the possibility that the neurons and other cells involved in the dopaminergic system can be sensitive to ketosis in a vertebrate with genes dysregulated in a similar manner (upregulation or downregulation) to patients with autism is extremely interesting, and it indicates that ketosis-induced adjustment of dysregulated genes may not be sufficient to mitigate cellular rhythms associated with insomnia.

Although this prediction, we do not consider our finding is directly applicable to human disorders without careful interpretation because systemic and organ physiologies are vastly different between fish and mammals. The knowledge gained from this unique fish system hinted a good starting point to investigate ketosis-induced behaviors in other asocial fish species and mammals.

This study, including the results of ketone bodies treatment, sheds light on the candidate genetic and molecular mechanisms associated with ketosis, deepening our knowledge of animal behaviors in response to metabolic states.

### The BHB supplement and body growth

In this study, ketone ester supplementation did not yield detectable negative impacts on growth, while the BHB sodium salt retarded body growth in surface fish. In contrast, the BHB salt treatment did not show a negative effect on cavefish growth (Additional file 12C, D and 15C, D). We suspect that the high level of sodium salt ingestion in the BHB salt treatment has a negative effect on growth, and the tolerance levels for high sodium ions (from BHB salt) may be different between surface fish and cavefish. Additionally, we also suspect that the reduction of nearby interaction in BHB-fed surface fish was caused by the high sodium ion levels (Additional file 14A). However, future physiological studies on the rhinal function in surface fish and cavefish are needed to provide answers.

### Ketosis in the cave environment

Cave-dwelling animals usually experience less temperature fluctuation and fewer dietary inputs [93], although these features can vary according to caves. The diets of cave-dwelling animals in the dry season (approximately 6 months/year) could consist of organic matter in pool bottoms, bat guano (for larger adults), or small crustaceans (for smaller fish), whereas food is sparse in the rainy season (approximately 6 months/year) [34, 94]. These available diets would contain extremely low amounts of carbohydrates and could be high in protein and fat (e.g., crustaceans). Although some amino acids, including lactate, and glycerol can be used for glucose synthesis in fish [95], cavefish are expected to be exposed to carbohydrate-deprived diets or frequent fasting and therefore experience frequent ketosis. In our observations, wild cavefish exhibited similar social affinity as observed in KD-fed cavefish in this study (Movie 1). Although these observations and dietary inputs suggest that wild cavefish may undergo frequent ketosis, recent multiple reports have indicated that cavefish may undergo anaerobic glycolysis to adapt to the cave water, which has approximately 20% lower oxygen levels (sevral cave pools [62, 96, 97]). Additionally, cavefish tend to store lipids instead of using them (via beta-oxidation) through the enhanced PPARγ pathway [36]. These expectations of low ketosis appear to contradict expectations in the wild—starved ketosis/aerobic conditions. However, they appears to align well with the findings in cavefish, namely higher blood glucose levels compared to surface fish in many feeding conditions (including KD feeding) in this study (Fig. 1, Additional files 4, 5 and 11). Cavefish appear to have evolved to maintain a high GKI (high blood glucose and low ketone levels); therefore, the physiology of cavefish may allow them to survive in low-oxygen conditions by utilizing anaerobic glycolysis. KD-fed cavefish behave similarly to wild cavefish because the balance between ketosis and glycolysis could reach a similar level as that in the wild after KD feeding. In contrast, if cavefish are fed a typical carbohydrate-rich lab fish diet, it may overactivate glycolysis and result in a higher GKI, which may lead to reduced social affinity and increased repetitive circling. The future use of a pharmacological glycolysis inhibitor (e.g., 2-deoxy-D-glucose; [98]) can reveal the relationship between GKI and cavefish behaviors.

### Summary statement

Surprisingly, solitary animals share a set of dysregulated genes and behavioral outputs (e.g., bees, and caveefish). In this study, we demonstrated that a diet inducing ketosis shifts these behaviors towards the surface fish phenotype, regardless of the presence of over a thousand dysregulated genes. There is a possibility that ketone body-based tratment, alongside ongoing gene therapy approaches, may open a path for sustainable and less toxic therapy for multigenic psychiatric disorders, including autism, although the affected gene pathways under ketosis remain unclear. Additionally, differentially expressed metabolic genes, which have been largely overlooked due to interpretational difficiulties, reemerged with their importance in understanding the genetics of behaivors. Given the highlighted role of mitochondria-based disorders in neuroscience [99, 100], investigating the balance between glycolysis and ketosis could serve as a starting point for identifying molecular mechanisms associated with neuronal states and behavioral shifts. Furthermore, reinvestigating the genetic factors for known evolved behaviors in the context of the metabolic shifts, in addition to the neural genes, may uncover the evolution of behaviors based on the evolution of metabolisms.

## Declarations

### Ethics approval and Consent to participate

The experimental protocols used in this study and fish care were approved by the institutional animal care and use committee (IACUC) at the University of Hawai‘i (17-2560).

### Consent to publish

All authors agreed on publishing these data and this manuscript.

### Availability of data and materials

The video datasets generated and/or analyzed during the current study are available at the university’s shared server and will be deposited to Zenodo (https://zenodo.org/). All program scripts used in this study are available at https://zenodo.org/record/5122894#.YvM4dPHMLsM

### Author contribution

MI: designed the experiments, performed the experiment and analyses, interpreted the data wrote the initial draft, and edited the manuscript.

AT: performed the BHB salt experiment and analysis, edited the manuscript

MG: performed the KE experiment and analysis, edited the manuscript

JC: performed the KE experiment and analysis, eidthed the manuscirpt

DB: performed the fasting experiment and analysis, edited the manuscript

VS: performed the BS vs CD experiment and analysis, edited the manuscript

MH: performed the fasting experiment and analysis, edited the manuscript

RBU: performed KD deprived exxperiment and analysis, edited the manuscript

KP: performed the blood measurements of the KE and BHB treatments analysis, edited the manuscript

MW: designed the experiments, consulted the experimental procedure, and edited the manuscript

RL: designed the experiments, consulted the experimental procedure, and edited the manuscript

MY: designed the experiments, performed the experiment and analyses, interpreted the data and wrote the initial draft, and edited the manuscript

## Acknowledgement

We thank C Balaan for constructive comments regarding insights on social-like interactions in cavefish. We are grateful to V Crystal, J Choi, L Lu, J Nguyen, VFL Fernandes, K Lactaoen, M Worsham, H Hernandez, N Doeden, J Kato, M Ito, E Doy, A Martinez, D Mones, H Yoshizawa for fish care assistance. We also thank to R Peres-David, B Yamamoto, B Suechting, AK Maunakea for their help in reading the UV absorption of the brain ketone bodies.

## Additional file, figure legends

**Additional file 1:**
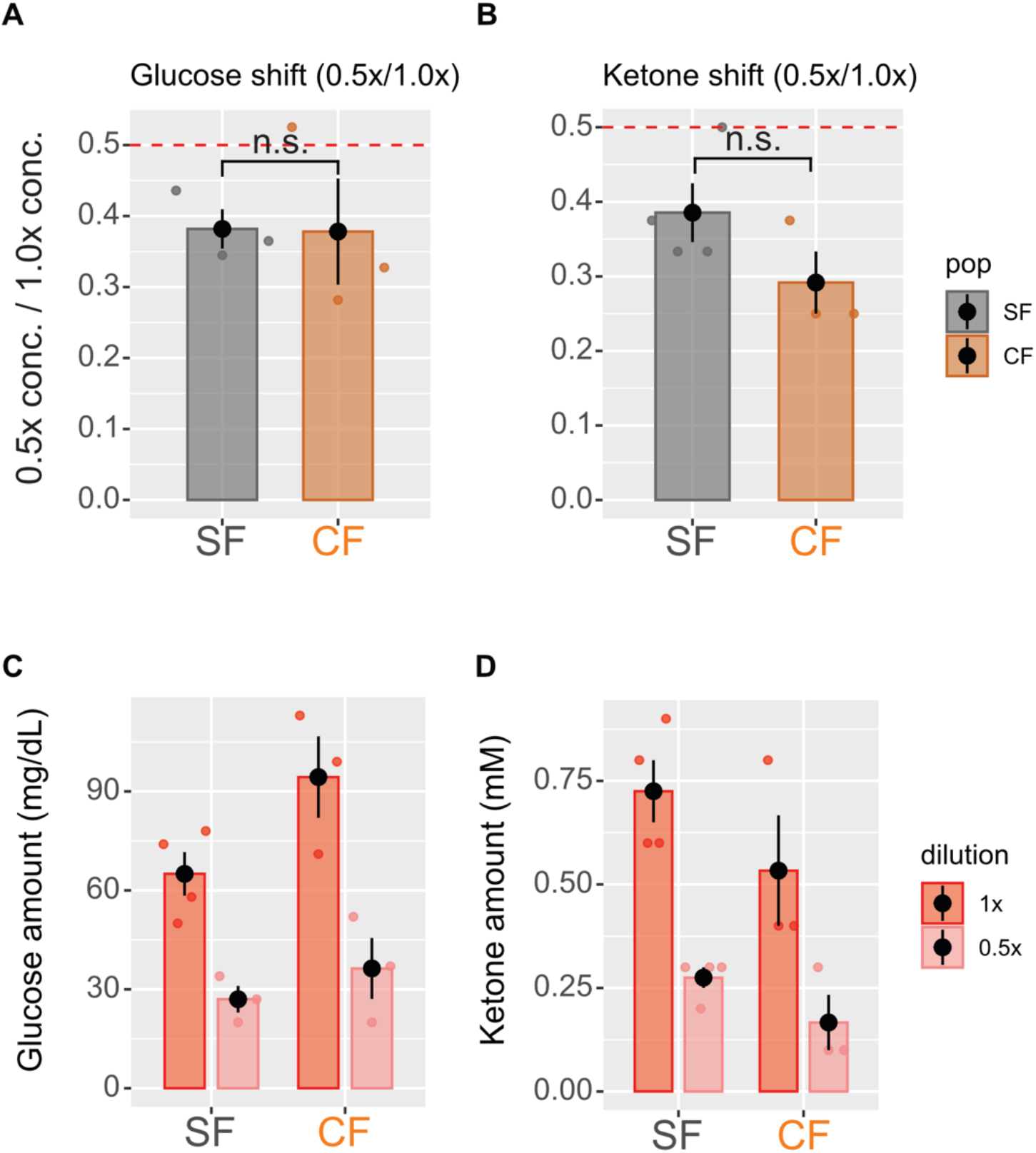
No detectable effect of the hematocrit levels on serum glucose or ketone body measurements. The blood of three surface fish and three cavefish was individually collected. Each blood sample was divided into non-diluted (1.0x) and diluted (0.5x) portions diluted with phosphate buffered saline (PBS). The blood samples were then measured using by Abbott Precision Xtra. If the difference of the hematocrit levels between surface and cavefish (28.51 ± 0.03% and 35.56± 0.03%, respectively) affected the readings of glucose or ketone bodies, the diluted/undiluted ratios would show a significant difference between surface fish and cavefish. (A) Ratio of glucose concentration in diluted (0.5x) and non-diluted (1.0x) blood. The expected ratio is 0.5, but it was shifted towards a lower concentration, possibly due to the low hematocrit level (∼15%) in 0.5x. However, the ratio of glucose concentrations showed no detectable difference between surface fish and cavefish, suggesting that the different hematocrit levels did not have a significant effect on the serum glucose readings. (B) Ratio of ketone bodies concentration in diluted (0.5x) and non-diluted (1.0x) blood. Similar to (A), the ratio showed no significant difference between surface fish and cavefish, indicating that the different hematocrit levels between the two populations had little effect on the ketone body concentration readings. (C) Raw measurements of the glucose concentrations used in (A). (D) Raw measurements of the ketone body concentrations used in (B). Detailed statistical scores are available in Additional file 2.

Additional file 2: **Statistical scores** for Figure 1-7; Additional file 1,4-15

https://docs.google.com/spreadsheets/d/1CFnKjQj2kQ1cvwpxl2N-Z6pEIOq69LtH/edit?usp=sharing&ouid=102214478617384425040&rtpof=true&sd=true

Additional file 3: **Movie file recorded in the Pachón cave pool in 2019**. Approximately 20 fish were transferred from the original cave pool to a foldable 2.44 x 2.44 m round pool (Play Day Round Kiddie Pool, Walmart Inc., Bentonville, Arkansas, USA) within a distance of approximately 10 m from the original pool in the same Pachón cave. These wild Pachón cavefish were acclimated for 1 day (24 hrs). Fish swimming behavior was then recorded using an infrared camcorder (DCR-SR200C, Sony, Tokyo, Japan)

https://drive.google.com/file/d/15iu09o_Jkl6vs1lwL0g52b5St8lZ5-LO/view?usp=sharing

**Additional file 4:**
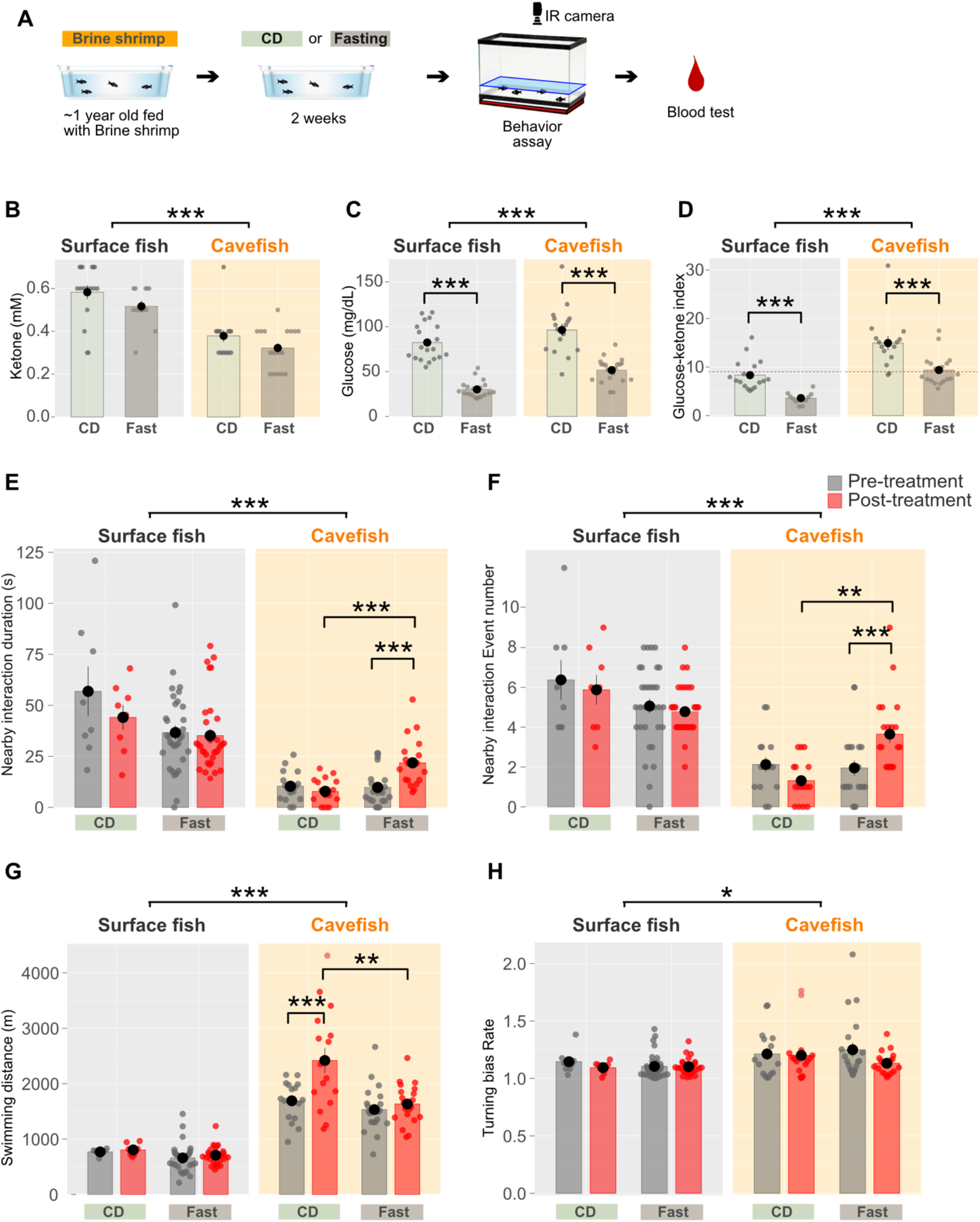
Two weeks of fasting reduced the glucose ketone index (GKI) in both surface fish and cavefish, and increased nearby interactions in cavefish. (A) Experimental procedure. Fish were either fasted (Fasting) or fed with brine shrimp (CD) for 2 weeks, after which blood glucose and ketone levels were measured. (B) Blood ketone level (mmol/L). The 2-week fasting did not result in detectable changes in ketone levels in either surface fish (SF) or cavefish (CF). Data are presented as the mean ± standard error of the mean. Dots indicate individual data points. (C) Blood glucose level (mg/dL). Fasting significantly reduced glucose levels in both SF and CF. (D) The glucose ketone index (GKI) indicated that the ratio of glucose to ketone was lowered by fasting in both SF and CF, suggesting that this diet altered the balance between glucose and ketone. (E) Duration of nearby interactions in the 5-min assays. The time periods when a fish was within proximity (≤5 cm) and spent more than 4 s in a 5-min assay were summed for each fish. Fasted cavefish showed longer nearby interactions than the fed control. (F) Number of nearby interaction events. The number of nearby events (≤5 cm and ≥4 s) in a 5-min assay was counted. Fasted cavefish showed a higher number of nearby interactions than the fed control. (G) Swimming distance in a 5-min assay. Cavefish swam a longer distance than surface fish, and the fasted cavefish swam a shorter than the CD-fed cavefish after the 2 weeks treatment. (H) Turning biases of surface fish (left) and cavefish (right). Cavefish exhibited a higher turning index than surface fish, but none of the treatments induced further turning bias. See also Figure 4. SF: N = 8 for CD feeding, N = 5 for fasting. CF: N = 8 for CD feeding, N = 8 for fasting. *: P < 0.05, **: P < 0.01, ***: P < 0.001. All detailed statistical data are available in Additional file 2.

**Additional file 5:**
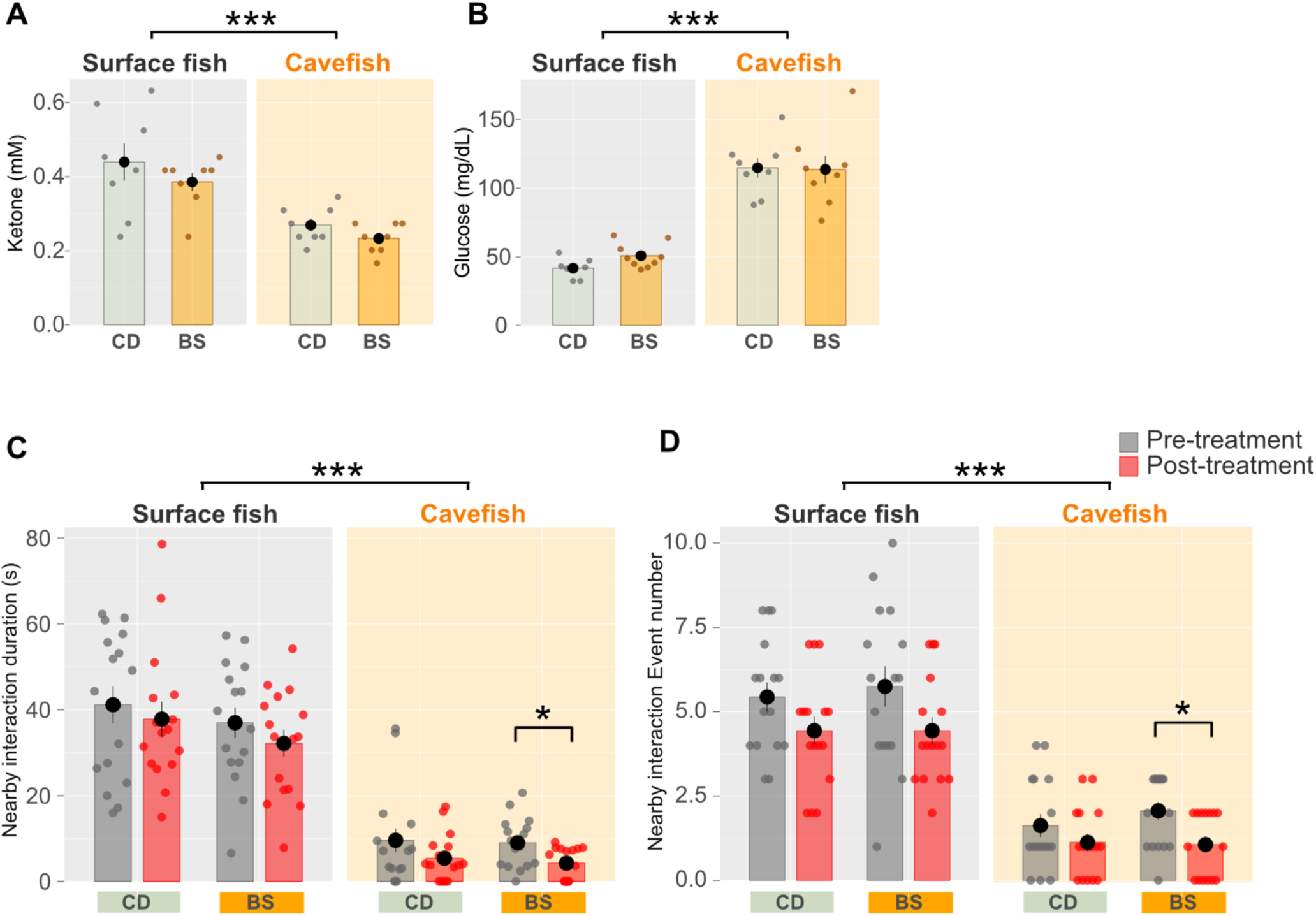
No detectable effect of the shift from live Artemia larvae (brine shrimp: BS) to the zebrafish standard control diet (CD) on the levels of serum glucose or ketone bodies, or nearby interactions. (A) Blood ketone level (mmol/L). After three weeks of treatment, ketone levels did not show detectable changes between CD and BS feeding in either surface fish or cavefish. Data are presented as the mean ± standard error of the mean. Dots indicate individual data points. (B) Blood glucose level (mg/dL). Glucose levels were not significantly changed by CD feeding in either surface or cavefish. (C) Duration of nearby interactions (s). During the 3 weeks of development, the average of the duration of nearby interactions was slightly reduced, but this was only detectable in BS-fed cavefish. (D) Number of nearby interaction events. The number of nearby interactions showed a similar result to (C): a slight reduction in interactions was detected in BS-fed cavefish. N = 20 for all groups. *: P < 0.05, **: P < 0.01, ***: P < 0.001. All detailed statistical data are available in Additional file 2. For (A) and (B), N = 8 for each group, For (C) and (D), N= 16 for each group. *: P < 0.05, **: P < 0.01, ***: P < 0.001. All detailed statistical data are available in Additional file 2.

**Additional file 6:**
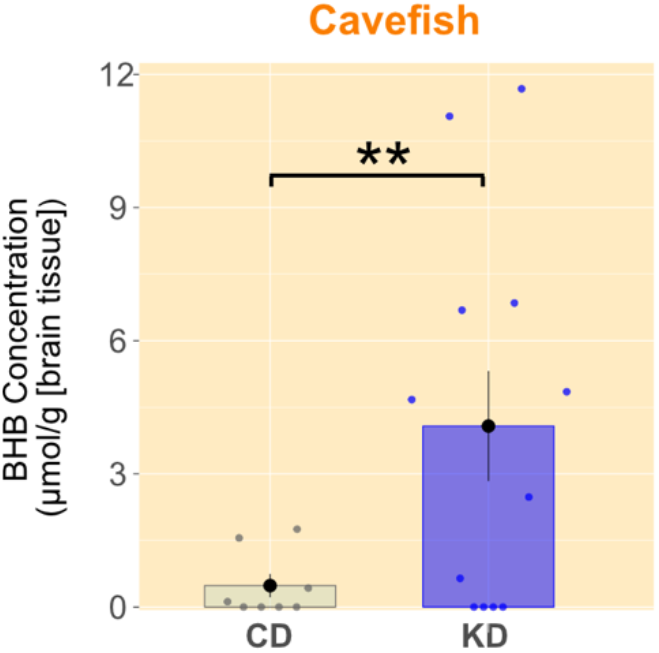
Brain ketone levels (beta-hydroxybutyrate) in cavefish under the CD or KD treatments. The concentrations of beta-hydroxybutyrate in the cavefish brain tissues were measured in 8 individuals fed the control diet (CD) and 12 individuals fed the ketogenic diet (KD) after 1 month of the diet treatment. The concentration was adjusted by the brain weight (g). Generalized linear model (family = Gamma) resulted in X^2^(1) = 10.5, P = 0.0012 **. N = 8 for the CD-fed cavefish, N = 12 for the KD-fed cavefish.

**Additional file 7:**
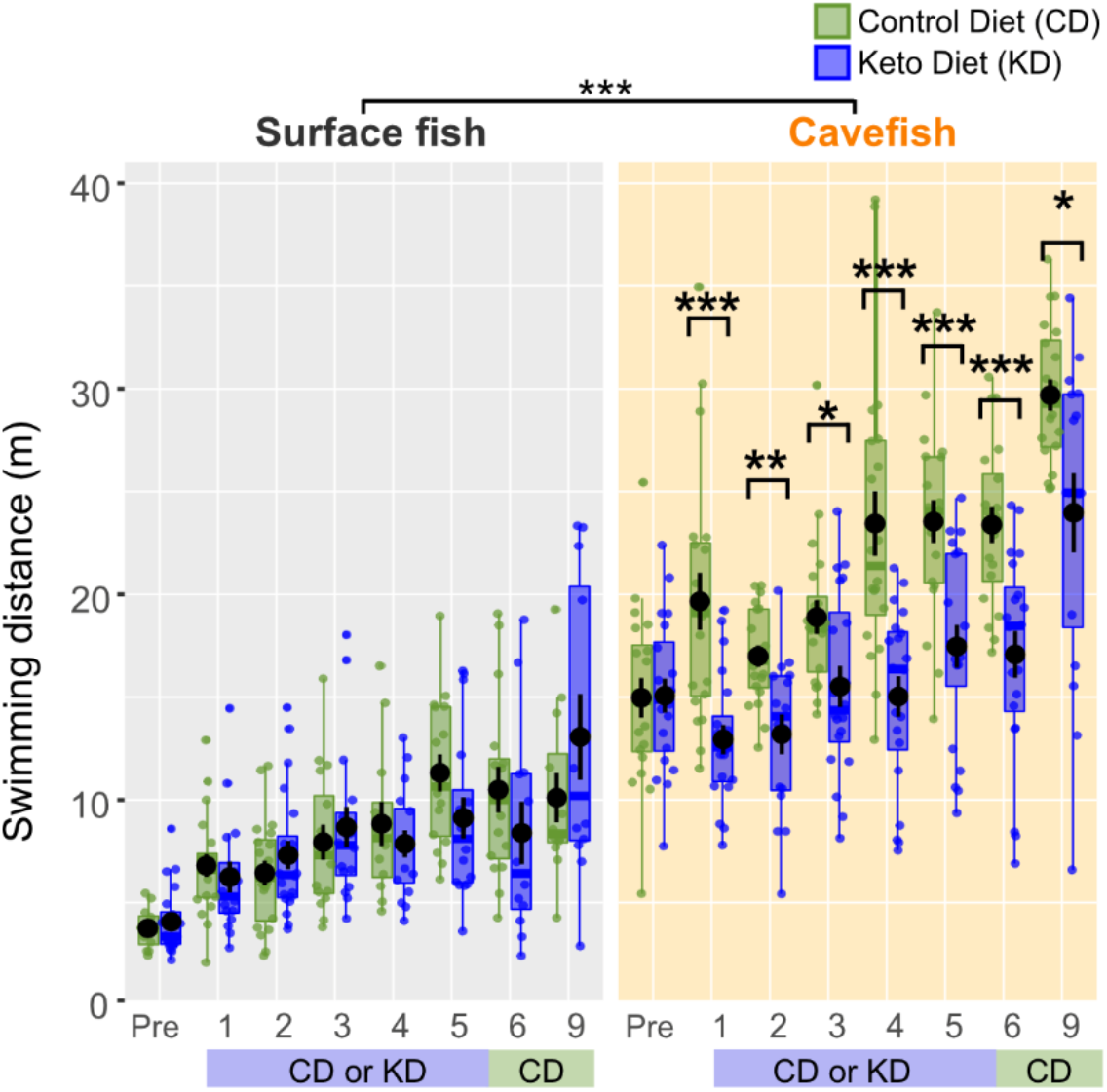
Ontogeny of swimming distance under ketogenic diet (KD) feeding. Surface fish exhibited an increase in the swimming distance over time under both the control diet (CD) and KD. In contrast, cavefish fed KD showed a suppression of the swimming distance and activity, starting in week 1, and these values remained smaller than those in CD-fed fish until week 9. However, KD-fed cavefish subsequently exhibited an increased swimming distance at week 9 after depriving KD feeding from week 6. Data are presented as box plots. Dots indicate individual data. N = 20 in all groups. *: P < 0.05, **: P < 0.01, ***: P < 0.001. All detailed statistical data are available in Additional file 2.

**Additional file 8:**
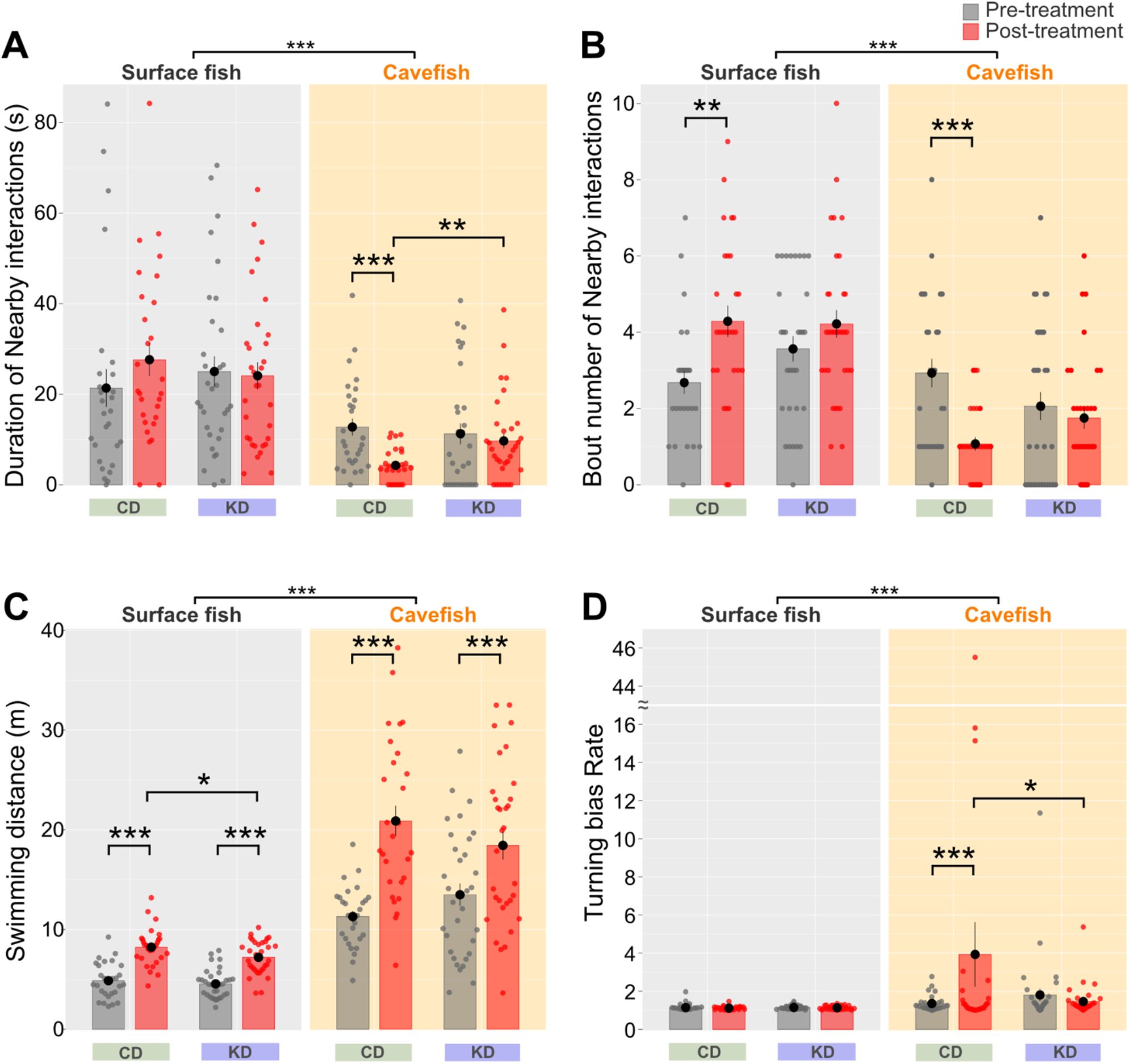
Consistent results were obtained in the repeated experiment for the duration and number of nearby interactions, and turning bias under control diet (CD) or ketogenic diet (KD) feeding. (A) Duration of nearby interactions, (B) Number of nearby interactions, (C) Swimming distance, and (D) Turning bias are presented. The overall tendencies were consistent with those observed in the original experiments, except for a shift in swimming distance (Figures 2 and 4). Surface fish did not exhibit any significant differences regarding the duration (A) or number of nearby interactions (B), the swimming distance (C), or turning bias (D). The duration (A) and number of nearby interactions (B) were maintained in KD-fed cavefish, while they were reduced in CD-fed cavefish, which also exhibited a higher level of turning bias (D). The swimming distance was not significantly reduced in KD-fed cavefish compared to CD-fed controls in this repeated experiment (C). Data are presented as the mean ± standard error of the mean. Dots indicate individual data. Surface fish: N = 28 for CD, N = 32 for KD. Cavefish: N = 28 for CD, N = 32 for KD. *: P < 0.05, **: P < 0.01, ***: P < 0.001. All detailed statistical data are available in Additional file 2.

**Additional file 9:**
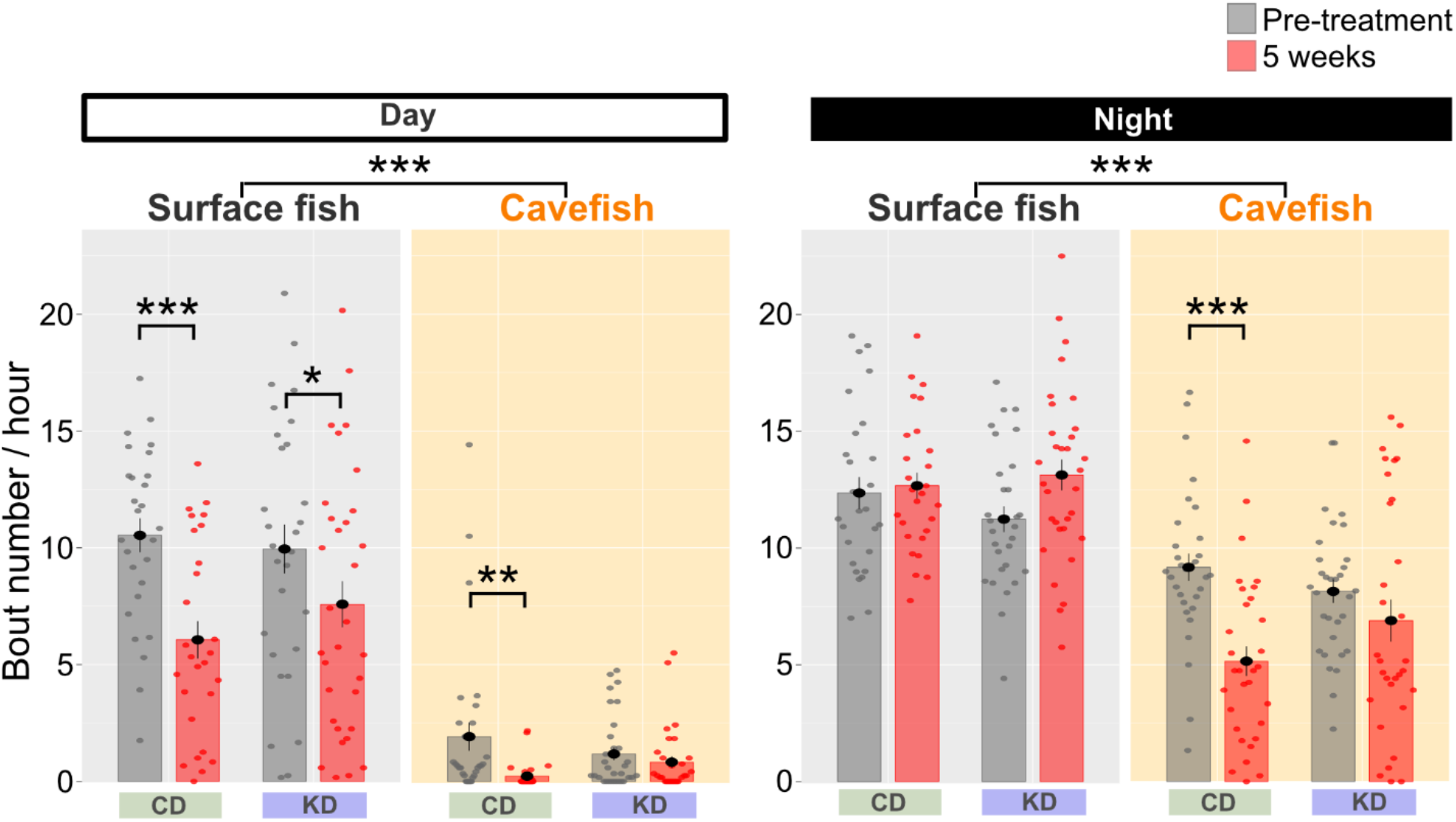
Daytime and nighttime number of sleeping events per hour under control diet (CD) or ketogenic diet (KD) feeding. After 5 weeks of growth, surface fish exhibited a reduced number of sleeping events during the day under both diets. CD-fed cavefish also exhibited a reduced number of sleeping events during the day and night. However, the number of sleeping events did not differ according to the diet in cavefish or surface fish. Data are presented as the mean ± standard error of the mean. Dots indicate individual data points. Surface fish: N = 28 for CD, N = 32 for KD. Cavefish: N = 28 for CD, N = 32 for KD. *: P < 0.05, **: P < 0.01, ***: P < 0.001. All detailed statistical data are available in Additional file 2.

**Additional file 10:**
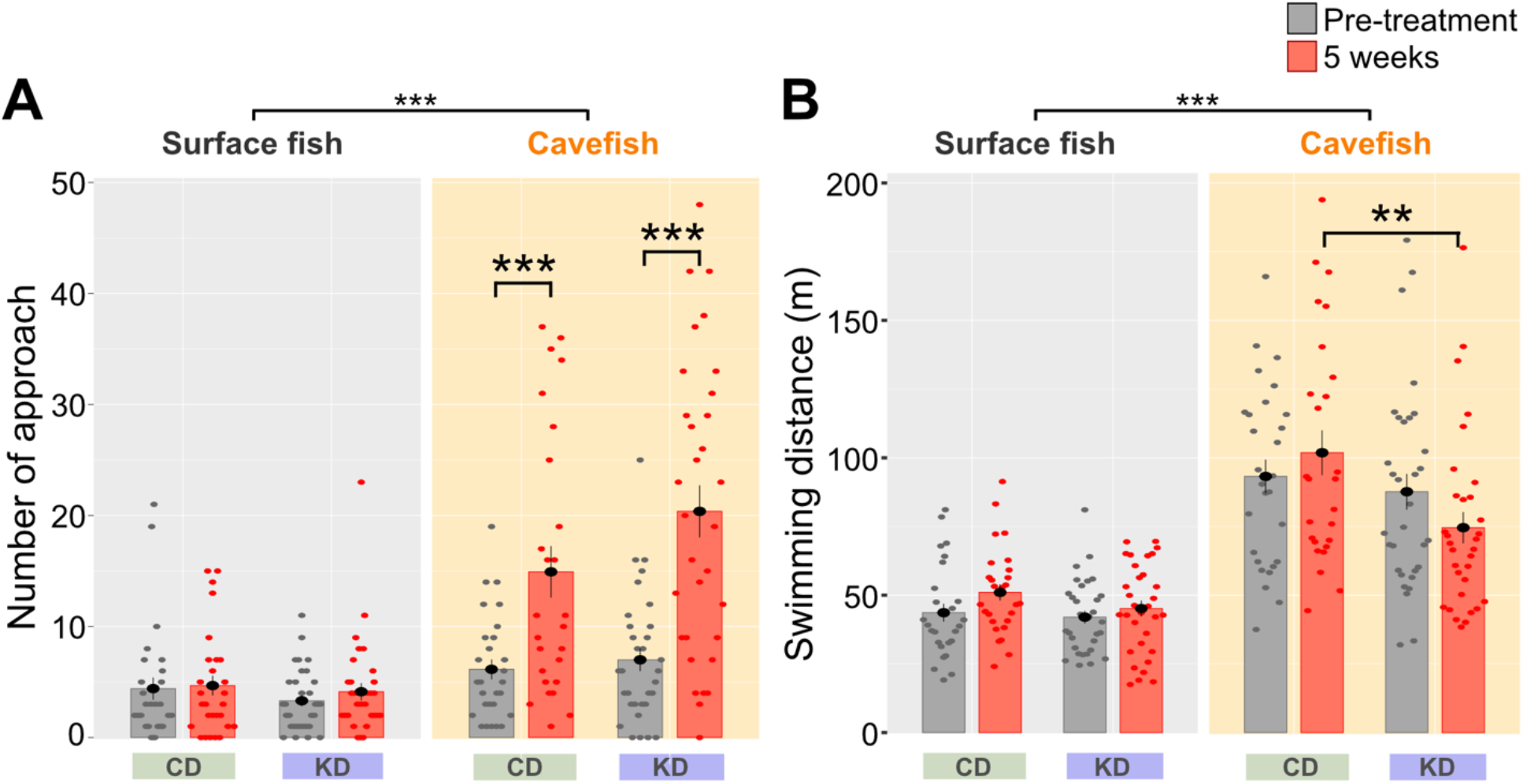
Vibration attraction behavior (VAB) and swimming distance during VAB under control diet (CD) or ketogenic diet (KD) feeding. (A) Number of approaches to the vibration rod in the 3-min assays. After 5 weeks of growth, the number of approaches increased in CD- and KD-fed cavefish, but no difference according to the diet was detected in either surface fish or cavefish. (B) Swimming distance during VAB. KD-fed cavefish swam significantly shorter distances than CD-fed cavefish, consistent with the nearby interaction and sleep studies. Data are presented as the mean ± standard error of the mean. Dots indicate individual data points. Surface fish: N = 28 for CD, N = 32 for KD. Cavefish: N = 28 for CD, N = 32 for KD. *: P < 0.05, **: P < 0.01, ***: P < 0.001. All detailed statistical data are available in Additional file 2.

**Additional file 11:**
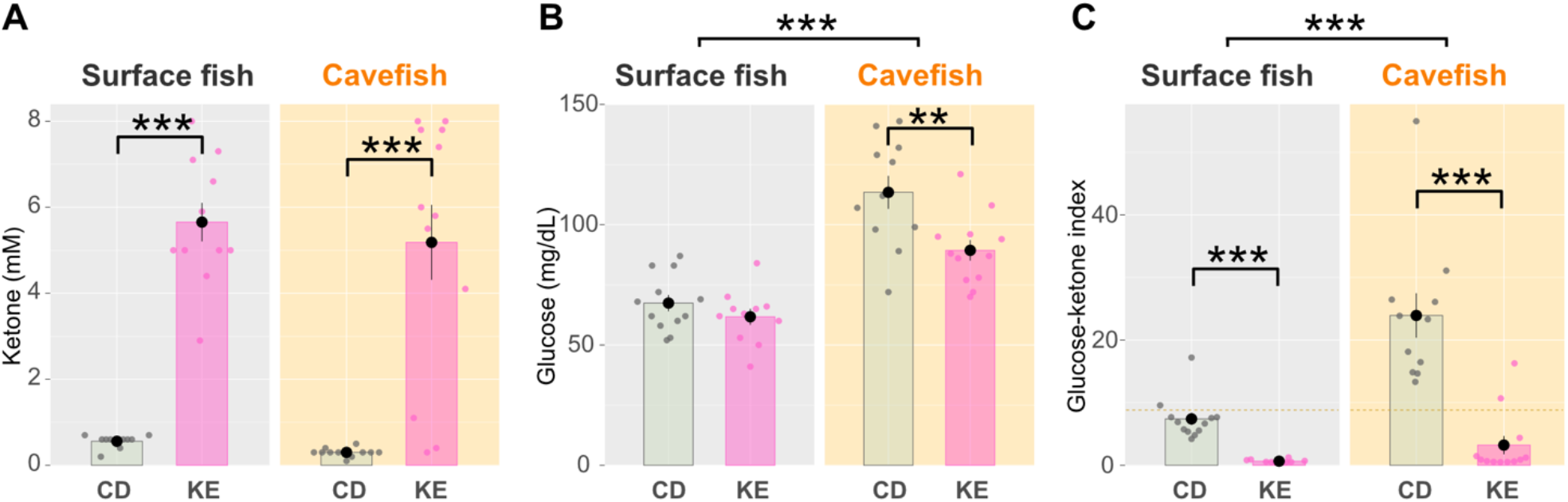
Five weeks of the ketone ester (KE)-supplemented diet feeding reduced glucose ketone index (GKI) in both surface fish and cavefish. (A) Blood ketone level (mmol/L). Ketone levels significantly changed after the 5-week feeding in both surface fish and cavefish. Data are presented as the mean ± standard error of the mean. Dots indicate individual data points. (B) Blood glucose level (mg/dL). Glucose levels were significantly reduced in cavefish (CF). (C) The glucose ketone index (GKI) indicated that the ratio of glucose to ketone was lowered by the 5-week KE supplemental feeding in both surface fish (SF) and CF, suggesting that this diet altered the balance between glucose and ketone. SF: N = 12 for CD feeding, N = 10 for KE feeding. CF: N = 11 for CD feeding, N = 12 for KE feeding. *: P < 0.05, **: P < 0.01, ***: P < 0.001. All detailed statistical data are available in Additional file 2.

**Additional file 12:**
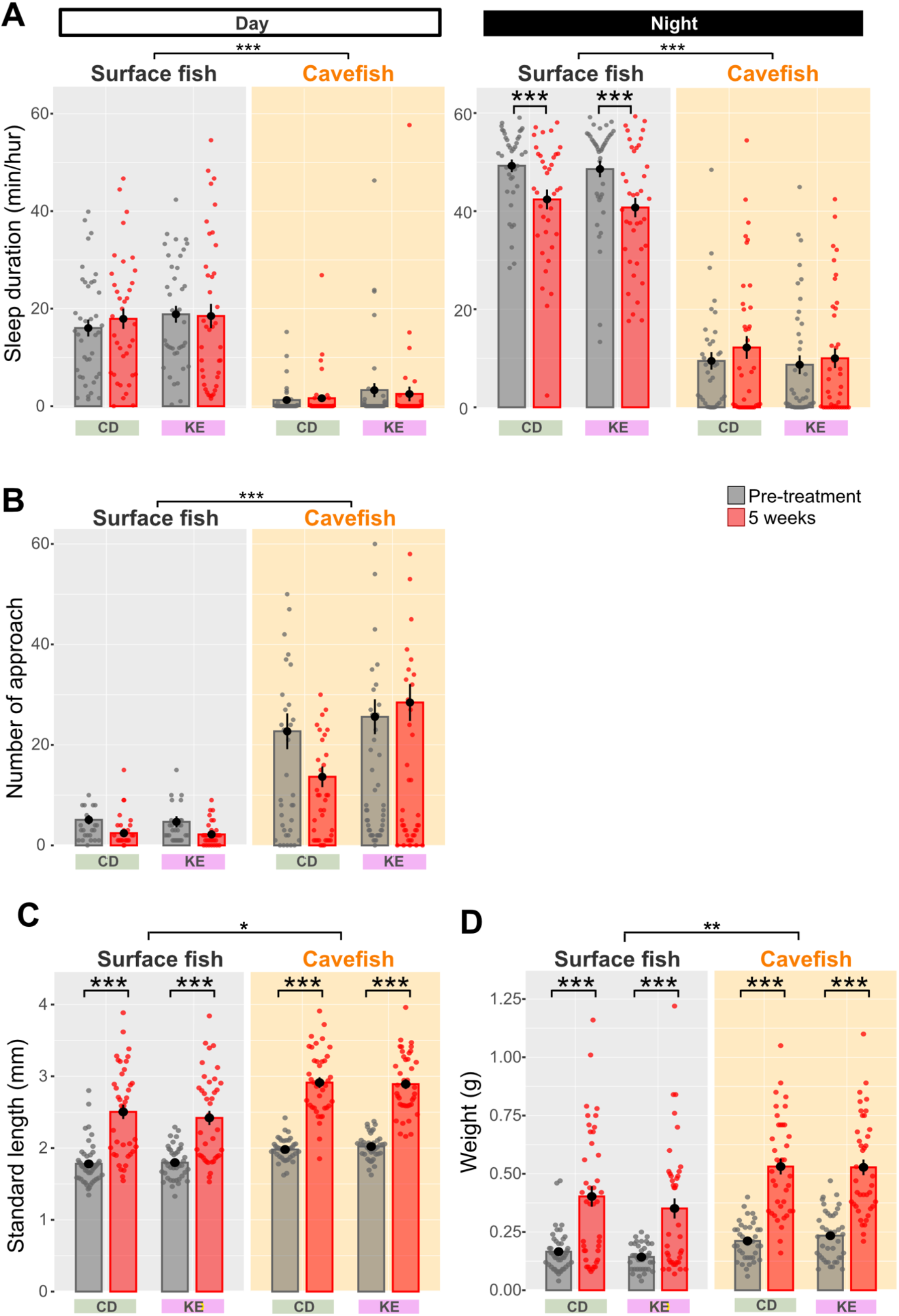
Day and night sleeping durations, vibration attraction behavior (VAB) and growth were not drastically changed by ketone ester-supplemented diet (KE) feeding. (A) Sleep duration (min/h) during the day (left) and night (right). During the 5 weeks of treatment, neither surface fish nor cavefish exhibited any detectable changes in KE feeding, although the night-time sleep in surface fish was reduced after 5 weeks of growth. (B) Number of approaches per the 3-min assay (VAB level). During the 5 weeks of treatment, the VAB level did not detectably shift in surface fish or cavefish regardless of diets. (C) Standard length (cm). KE-fed surface fish and cavefish were comparable to their CD-fed counterparts. (D) Body weight (g). KE-fed surface fish and cavefish weighed comparable to their CD-fed counterparts. Data are presented as the mean ± standard error of the mean. Dots indicate individual data points. For (A) and (B), Surface fish: N = 37 for CD, N = 39 for KE. Cavefish: N = 39 for CD, N = 40 for KE. For (C) and (D), Surface fish: N = 40 for CD, N = 39 for KE. Cavefish: N = 39 for CD, N = 40 for KE. *: P < 0.05, **: P < 0.01, ***: P < 0.001. All detailed statistical data are available in Additional file 2.

**Additional file 13:**
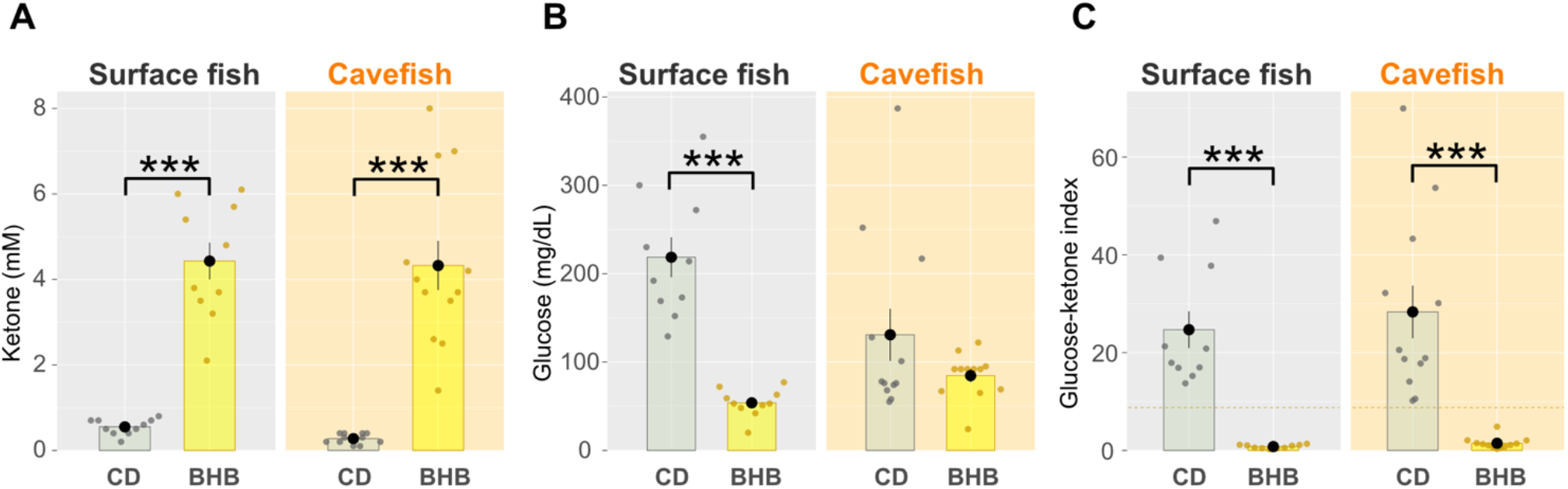
Four weeks of the BHB salt-supplemented diet feeding reduced glucose ketone index (GKI) in both surface fish and cavefish. (A) Blood ketone level (mmol/L). Ketone levels significantly changed by the 4-week feeding in both surface fish (SF) and cavefish (CF). Data are presented as the mean ± standard error of the mean. Dots indicate individual data. (B) Blood glucose level (mg/dL). Glucose levels were significantly reduced in SF. (C) The glucose ketone index (GKI) indicated that the ratio of glucose to ketone was lowered by the 4-week BHB salt-supplemented feeding in both SF and CF, suggesting that this diet altered the balance between glucose and ketone. SF: N = 10 for CD feeding, N = 10 for BHB feeding. CF: N = 10 for CD feeding, N = 10 for BHB feeding. *: P < 0.05, **: P < 0.01, ***: P < 0.001. All detailed statistical data are available in Additional file 2.

**Additional file 14:**
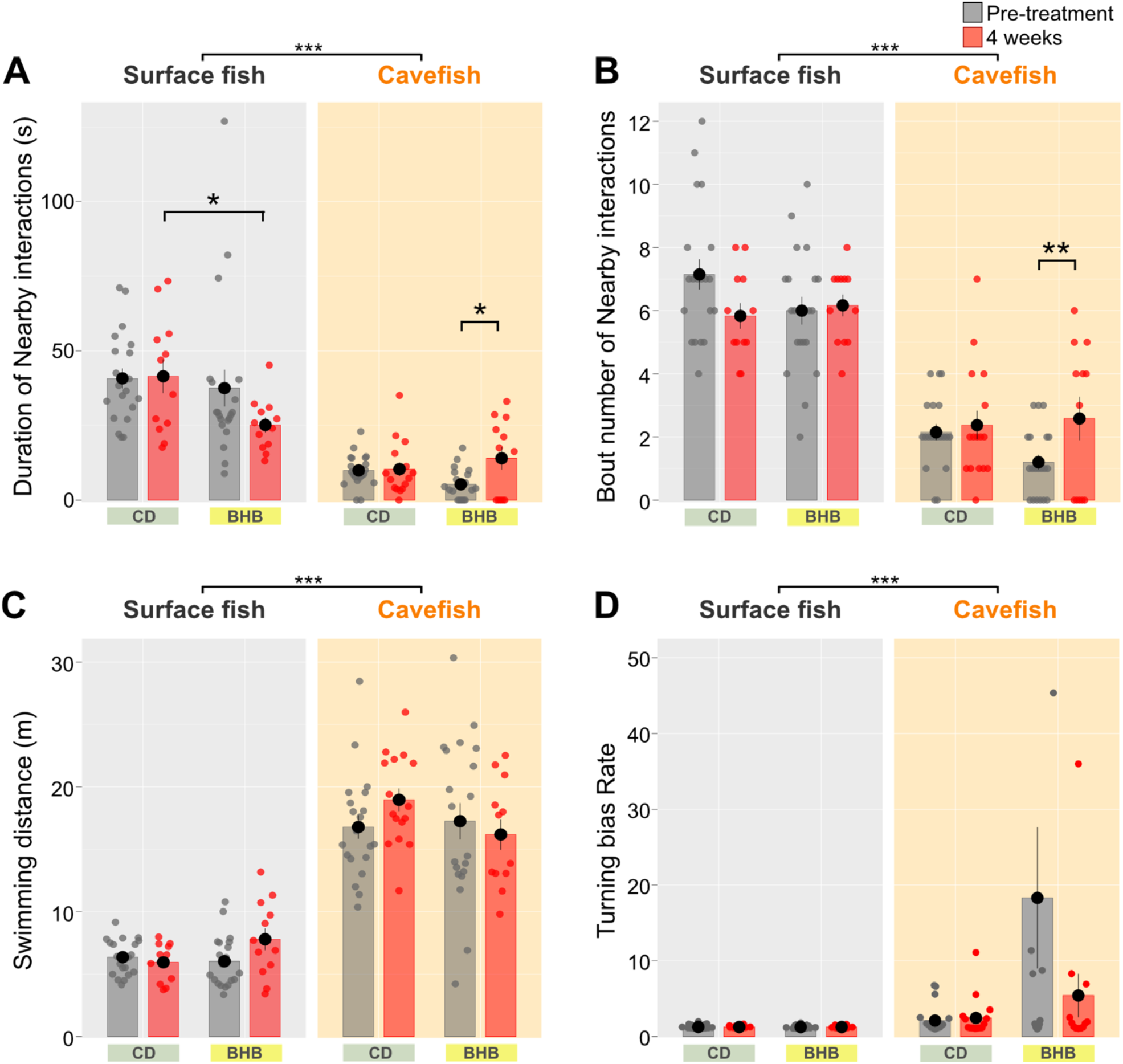
Nearby interactions and other behaviors under control diet (CD) or BHB sodium salt–supplemented diet (BHB) feeding. (A) Duration of nearby interactions (s). After 4 weeks, the duration of nearby interactions decreased in BHB-treated surface fish and increased in BHB-treated cavefish. (B) Number of nearby interactions. The number of nearby interactions increased in BHB-treated cavefish. (C) Swimming distance. No difference was detected between the CD and BHB groups. (D) Turning bias ratio. BHB-treated cavefish tended to exhibit decreased biased turning, although this reduction was not significant. Data are presented as the mean ± standard error of the mean. Dots indicate individual data points. N = 20 for all groups. *: P < 0.05, **: P < 0.01, ***: P < 0.001. All detailed statistical data are available in Additional file 2.

**Additional file 15:**
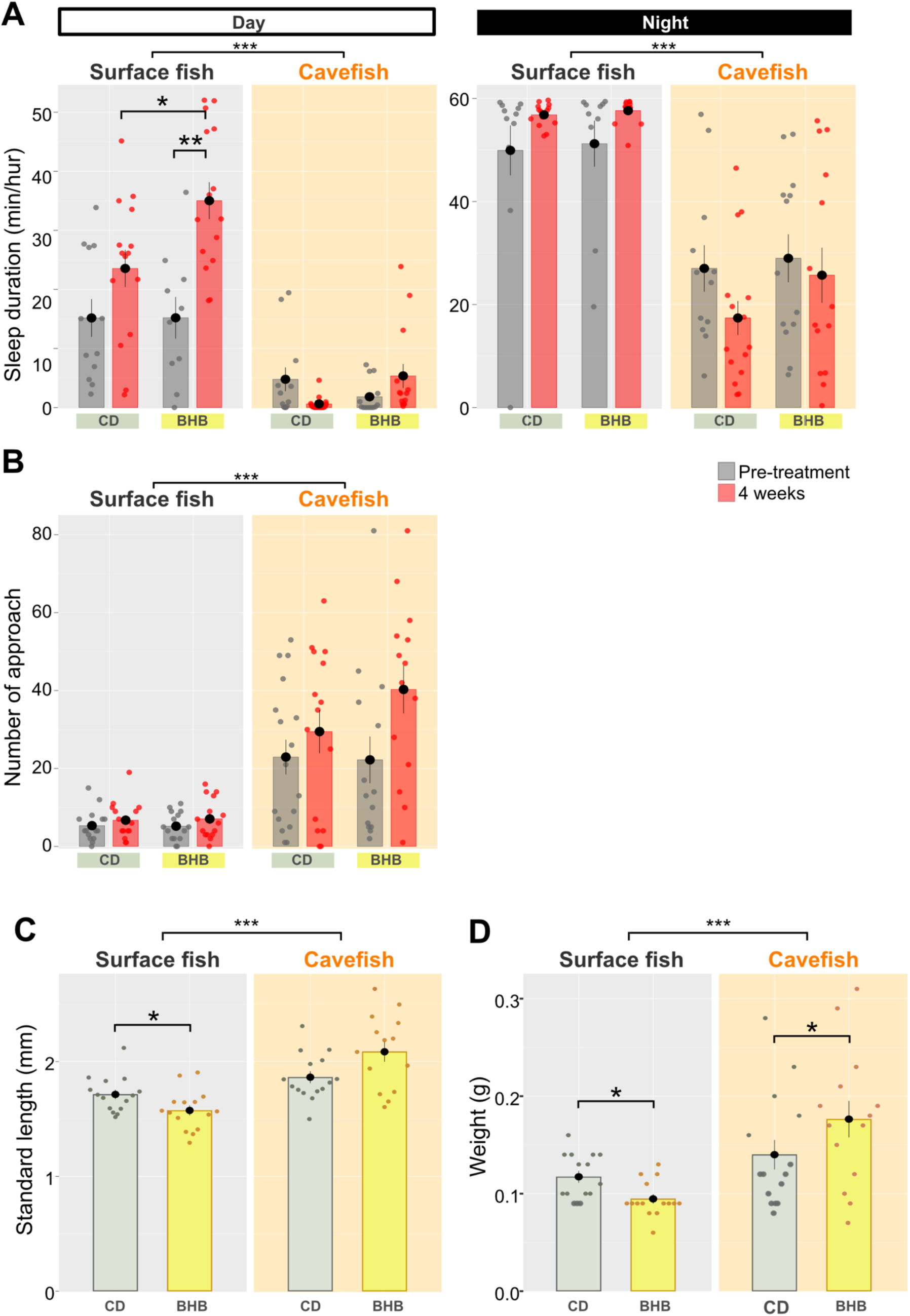
Day and night sleeping durations and vibration attraction behavior (VAB) were not drastically changed by beta-hydroxybutyrate-supplemented diet (BHB) feeding. (A) Sleep duration (min/h) during the day (left) and night (right). During 4 weeks of treatment, the daytime sleep duration increased in surface fish under BHB feeding but not during nighttime. Cavefish did not exhibit any detectable changes. (B) Number of approaches per the 3-min assay (VAB level). During 4 weeks of treatment, the VAB level did not shift in surface fish or cavefish regardless of the diets. Data are presented as the mean ± standard error of the mean. Dots indicate individual data points. N = 20 for all groups. (C) Standard length (cm). The BHB-treated surface fish grew slower than the control diet-treated ones. Cavefish showed no detectable change between the control and BHB supplementation. (D) Body weight (g). BHB-treated surface fish exhibited significantly reduced weight, whereas the weight of BHB-fed cavefish did not show any negative effect on the growth. Data are presented as the mean ± standard error of the mean. Dots indicate individual data. N = 20 for all groups. *: P < 0.05, **: P < 0.01, ***: P < 0.001. All detailed statistical data are available in Additional file 2.

Additional file 16: **Table of possible biological processes in each behavior tested in this study**

https://drive.google.com/file/d/1LdXTn68vlqU7I-bxra6k5TCNj4-NcGdz/view?usp=sharing

Additional file 17: **GO term re-analysis of the 72-hour post-fertilization transcriptome**

For the RNAseq transcriptome analysis, variation in gene expression was analyzed using previously published RNAseq data (GenBank Sequence Read Archive (SRA), accession code: PRJNA258661) [26, 92, 101]. The data were analyzed by following previously published protocols [31, 102]. Gene ontology terms (GO terms) were analyzed using the AmiGO 2 platform [103].

https://docs.google.com/spreadsheets/d/1Lh2Iegpa-kikYvU8BCS9ZMM95oLwReOp/edit?usp=sharing&ouid=102214478617384425040&rtpof=true&sd=true

Additional file 18: **KEGG pathway analysis used the data in Yoshizawa et al., 2018**

https://drive.google.com/file/d/1LgI201j4d2R7FIwOC5Z7BlMQLVVwgkNU/view?usp=sharing

